# Cleavage of the RNA polymerase II general transcription factor TFIIB tunes transcription during stress

**DOI:** 10.1101/2025.07.28.667251

**Authors:** Leah Gulyas, Azra Lari, Sahil B. Shah, Britt A. Glaunsinger

**Affiliations:** Department of Plant and Microbial Biology, University of California, Berkeley, CA 94720, USA; Center for Computational Biology, University of California, Berkeley, Berkeley, CA, USA; Department of Molecular and Cell Biology, University of California, Berkeley, CA 94720, USA; Howard Hughes Medical Institute, University of California, Berkeley, CA 94709, USA

**Author notes:** Authors contributed equally to this work (Azra Lari, Sahil B. Shah).

## Abstract

Cellular stressors often cause widespread repression of RNA polymerase II (RNAP II) activity, which is thought to facilitate a focused transcriptional output towards stress resolution. In many cases, however, the underlying regulatory mechanisms remain unknown. Here, we demonstrate that stress-induced downregulation of the general transcription factor TFIIB tempers expression of specific stimulus response genes. Following a variety of stressors, TFIIB is proteolytically cleaved between its cyclin folds at conserved aspartic acid residue D207 by caspases- 3 and 7. Cleavage in this portion of the protein significantly reduces the ability of TFIIB to form a TBP-TFIIB-DNA promoter complex in vitro. Using both overexpression and endogenous base-editing, we find that B and T cells that are unable to cleave TFIIB upregulate expression of a select gene set during apoptosis. These TFIIB-sensitive genes are primarily short, stimulus-responsive and proto-oncogenic loci, and cleavage of TFIIB temporally restricts their expression. Failure to cleave TFIIB during stress leads to aberrant lymphocyte proliferation during chemical perturbation. Hence, caspase targeting of TFIIB destabilizes transcription to tune gene expression, allowing for proper stress resolution.

## Introduction

Cell stress is accompanied by rapid changes to the transcriptome. This involves the induction of genes required for stress resolution, but is also frequently associated with broad downregulation of basal gene expression by RNA polymerase II (RNAP II) (Gulyas and Glaunsinger 2022). Suppression of RNAP II activity occurs in response to various stressors, including DNA damage (Tufegdžić Vidaković et al. 2020), hyperosmotic stress (Rosa-Mercado et al. 2021), pro-apoptotic signals (Duncan-Lewis et al. 2021), and viral infection (Abernathy et al. 2015; Bauer et al. 2018). Transcriptional repression may focus gene expression on the stress response by freeing resources, as well as restricting viral access to essential transcription machinery and/or permitting cells to properly execute cell death pathways. While some drivers of stress-induced transcriptional repression have been well studied, in many cases the underlying mechanisms remain unknown. We previously found that transcriptional repression in gammaherpesvirus-infected cells coincides with a reduction in the protein abundance of some RNAP II machinery, most notably the general transcription factor TFIIB (Hartenian et al. 2020).

TFIIB is crucial for RNAP II recruitment and initiation. It stabilizes the general transcription factor TFIID on promoter DNA, and its N-terminal B-ribbon threads to the RNAP II core complex, drawing it to the promoter (Deng and Roberts 2007; Kostrewa et al. 2009). TFIIB promoter binding is transient, with rapid dissociation and reassociation observed *in vitro* (Zhang et al. 2016). Once stably bound, TFIIB assists RNAP II in unwinding DNA and identifying the transcription start site (Kostrewa et al. 2009; Sainsbury et al. 2013). TFIIB also plays a more enigmatic role in 3′ transcript processing in yeast (Singh and Hampsey 2007; O’Brien and Ansari 2022; O’Brien and Ansari 2023), as well as in transcription termination in yeast and mammalian cells (Wang et al. 2010; Santana et al. 2022; O’Brien et al. 2024).

Acute global loss of TFIIB broadly represses gene expression, confirming its central role in cellular transcription (Santana et al. 2022). However, several observations suggest that it may be a point of control during stress. For example, its abundance fluctuates during differentiation (Shiraishi et al. 2009) and retinal damage (Sang et al. 2013), it is depleted during murine gammaherpesvirus infection as mentioned above (Hartenian et al. 2020), and it is relocalized to the cytoplasm by Thogoto virus to alter immune gene expression (Haas et al. 2018). TFIIB is further subject to regulation through protein modifications including phosphorylation (Wang et al. 2010; Shandilya et al. 2012), acetylation (Choi et al. 2003), and ubiquitylation (Watanabe et al. 2020).

Here, we reveal that a variety of stresses regulate TFIIB abundance via a caspase-mediated cleavage event in B and T lymphocytes. This occurs at a conserved residue located between the two cyclin folds of TFIIB that contact promoter DNA. In vitro, cleavage in this region decreases TFIIB’s ability to bind to DNA. We used exogenous expression in B cells and endogenous base editing in T cells to show that an uncleavable allele of TFIIB primarily upregulates a subset of short, stimulus-responsive genes induced during apoptosis. Cells that cannot cleave TFIIB to dampen expression of these genes aberrantly proliferate during chemical stress. Together, our data suggest that TFIIB cleavage tempers transcription at stress response and mitogenic genes in apoptotic lymphocytes, functioning as a stress-activated mechanism to drive transcriptional repression.

## Results

### Cellular stress induces caspase cleavage of TFIIB at residue D207

To explore whether TFIIB protein abundance is regulated during stress, we focused on immune cells, as B and T lymphocytes are highly sensitive to stress (Muralidharan and Mandrekar 2013). B cells are also a primary target of several gammaherpesviruses, which have been previously shown to downregulate TFIIB during viral lytic replication (Hartenian et al. 2020). We first assessed TFIIB protein levels in a B cell lymphoma line (TRExRTA BCBL-1, henceforth referred to as BCBL-1) that harbors Kaposi’s sarcoma-associated herpesvirus (KSHV) in a latent state (Nakamura et al. 2003). The cells were subjected to several stressors, including the DNA damage agent etoposide, the translation inhibitor cycloheximide, or lytic reactivation of KSHV via doxycycline (dox)-induced expression of the master viral lytic activator, RTA. Western blotting revealed that all three treatments reduced endogenous full-length TFIIB protein relative to its levels in control cells, suggesting that it is commonly targeted during stress **(Fig. 1A)**. Notably, the reduction in full-length TFIIB was accompanied by the appearance of smaller molecular weight bands in the ∼20 kD range. We detected a similar cleavage event in cells stably expressing a 2x C-terminal strep-tagged version of TFIIB, which yielded a ∼14 kD strep-tagged product, suggesting that TFIIB proteolysis occurs closer to the C-terminus of the protein **(Fig. 1A)**.

**Figure 1.**
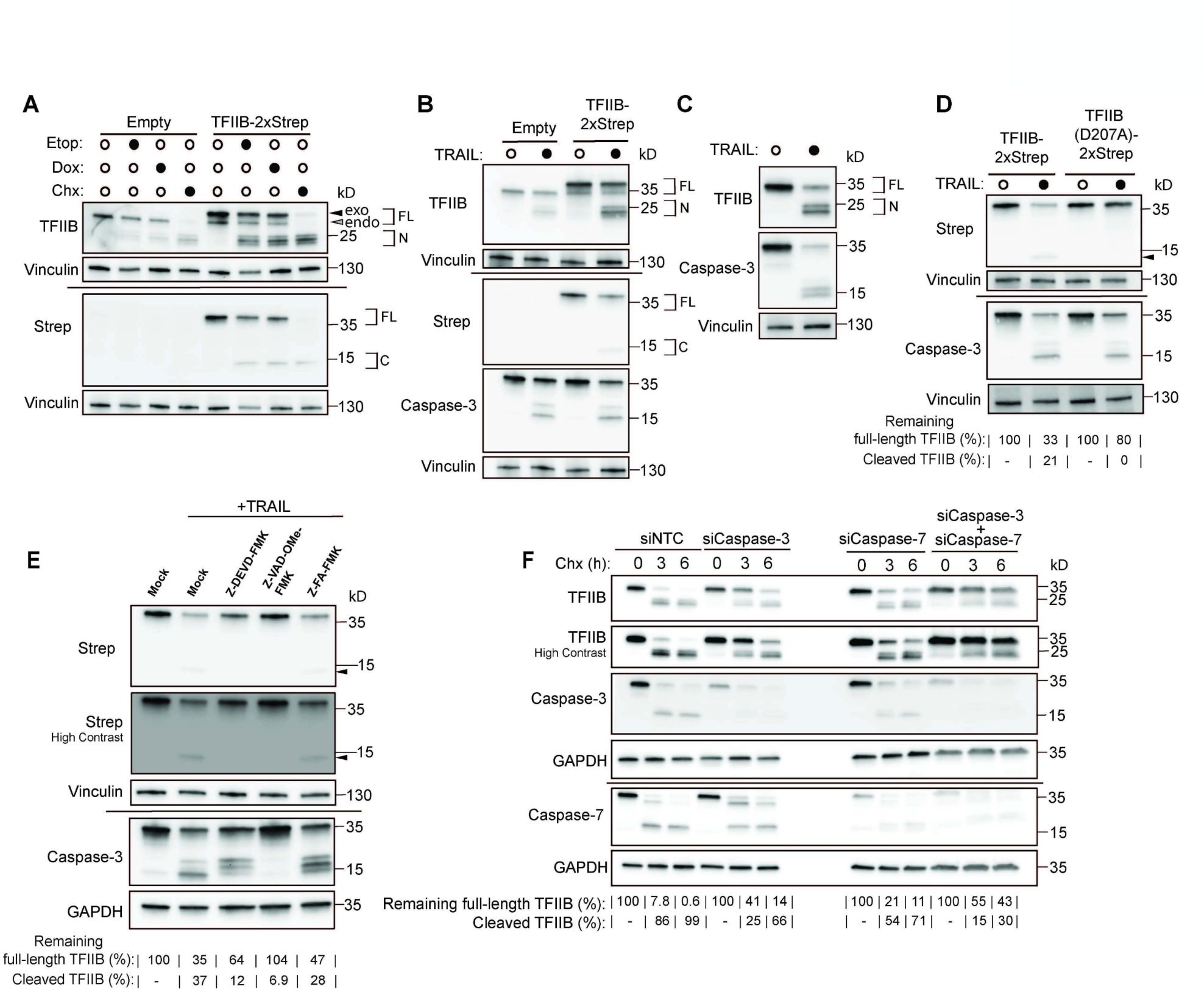
Cellular stress depletes TFIIB via caspase-3/7 cleavage at the D207 residue. **(A)** BCBL-1 cells expressing empty vector or TFIIB-2xStrep were mock-treated or treated with 25 μM etoposide (Etop) or 2 μg/mL doxycycline (Dox) for 24h, or 100 μg/mL cycloheximide (Chx) for 6 h and analyzed by western blotting. Black arrow indicates full length strep-tagged TFIIB and gray arrow denotes full-length endogenous TFIIB. FL = full-length protein, N = N-terminal cleavage product, C = C-terminal strep-tagged cleavage product for *A-C*. For this and subsequent blots, vinculin or GAPDH is a loading control and anti-TFIIB recognizes both endogenous and strep-tagged constructs. **(B)** Western blot of BCBL-1 cells expressing the indicated constructs and treated +/- 100 ng/mL TRAIL ligand for 6 h. Caspase-3 cleavage is indicative of its activation. **(C)** Western blot of Jurkat cells +/- 166 ng/mL TRAIL for 6 h. **(D)** As in *(B)* for 8 h. Black arrow indicates C-terminal cleavage product for *D-E.* **(E)** Western blot of BCBL-1 cells expressing TFIIB-2xStrep and treated with 80 μM of the indicated inhibitor for 2 h, followed by treatment +/- 500 ng/mL TRAIL ligand for 7 h. Contrast is enhanced as indicated for visualization of the cleavage product. **(F)** BCLB-1 cells were nucleofected with the indicated control (siNTC) or targeting siRNAs for 48 h followed by treatment with 100 μg/mL Chx for 0, 3, or 6 h and western blotting. For quantifying the impact on TFIIB cleavage for *D-F*, remaining full-length TFIIB is the percentage of load-normalized full-length protein relative to the 0 h or mock timepoint for each condition. Cleaved TFIIB is the percentage of cleavage product relative to total TFIIB protein in each lane.

To explore whether cleavage might preferentially occur on the transcriptionally active or inactive form of TFIIB, we individually mutated two well-characterized TFIIB regulatory residues, S65 and R66. Residue S65 undergoes phosphorylation to promote transcription at a subset of genes (Shandilya et al. 2012), and the R66E mutation stabilizes an inhibitory “closed” TFIIB conformation (Hawkes et al. 2000; Glossop 2004). However, neither residue influenced TFIIB cleavage, as phosphomimetic S65E, unphosphorylatable S65A TFIIB, and the R66E mutant all showed the same cleavage pattern as the wild-type (WT) **(Supplemental Fig. S1A-B)**. While other residues or modifications may regulate TFIIB cleavage, these data suggest that cleavage may occur independently of its transcriptional activity.

Stress-induced proteolytic cleavage is a hallmark of caspase activity. The addition of the apoptosis-inducing TRAIL ligand, which activates the executioner procaspase-3, to BCBL-1 cells and to Jurkat T cells also resulted in TFIIB cleavage **(Fig. 1B-C)**. We identified a putative caspase-3 cleavage site after TFIIB aspartic acid residue 207 using CaspSites, a repository for caspase substrates (Wang and Julien 2023). Mutation of this site to alanine (D207A) in strep-tagged TFIIB rendered the protein resistant to cleavage, confirming this as the primary cleavage site **(Fig. 1D)**. Both the pan-caspase inhibitor Z-VAD-OMe-FMK and the caspase-3/7 inhibitor Z-DEVD-FMK impaired TFIIB-2xStrep cleavage during TRAIL-induced apoptosis relative to the Z-FA-FMK negative control **(Fig. 1E)**. Caspase-3 and caspase-7 share overlapping substrate motifs, and we thus assessed their relative contribution to TFIIB cleavage (McStay et al. 2008; Julien and Wells 2017). Individual siRNA knockdown of caspase-3 or caspase-7 extended the half-life of full-length TFIIB, with a more pronounced effect observed for caspase-3 and the strongest effect upon co-depletion of both caspases **(Fig. 1F)**. Thus, TFIIB is cleaved by caspases-3 and 7 at residue D207, decreasing its intact protein levels during a variety of cellular stresses.

### TFIIB cleavage impairs promoter binding in vitro

We next examined the implications of TFIIB cleavage at D207. Sequences surrounding the TFIIB caspase cleavage site are highly conserved across metazoans, and the D207 residue is maintained in fruit flies through humans, although not in yeast, hydra, or nematodes **(Fig. 2A)**. This may suggest an early evolutionary advantage in controlling TFIIB protein abundance as increasingly complex apoptotic programs developed. D207 is located in a short, unstructured region that bridges the two positively charged cyclin folds of TFIIB, which contact promoter DNA **(Fig. 2B)**, suggesting that the separation of these domains could compromise TFIIB’s ability to bind to the promoter. To test this hypothesis, we purified a version of TFIIB containing a PreScission protease site immediately after the D207 residue (TFIIB^PS^) to induce cleavage in vitro. After confirming that TFIIB^PS^ could be effectively proteolyzed by PreScission **(Supplemental Fig. S2A)**, TFIIB^PS^ promoter binding was assessed by an electrophoretic mobility shift assay (EMSA). In contrast to full-length TFIIB^PS^, which shifted Adenovirus Major Late (AdML) promoter DNA in the presence of TATA-binding protein (TBP), cleavage of TFIIB before or during the reaction significantly impaired the shift and resulted in a greater pool of free probe **(Fig. 2C)**. This was not due to simple steric hindrance as the binding capacity of TFIIB lacking the engineered protease cleavage site was unaffected by PreScission **(Supplemental Fig. S2B)**. Thus, TFIIB cleavage in this region of the protein impairs its ability to stably bind promoter DNA in vitro.

**Figure 2.**
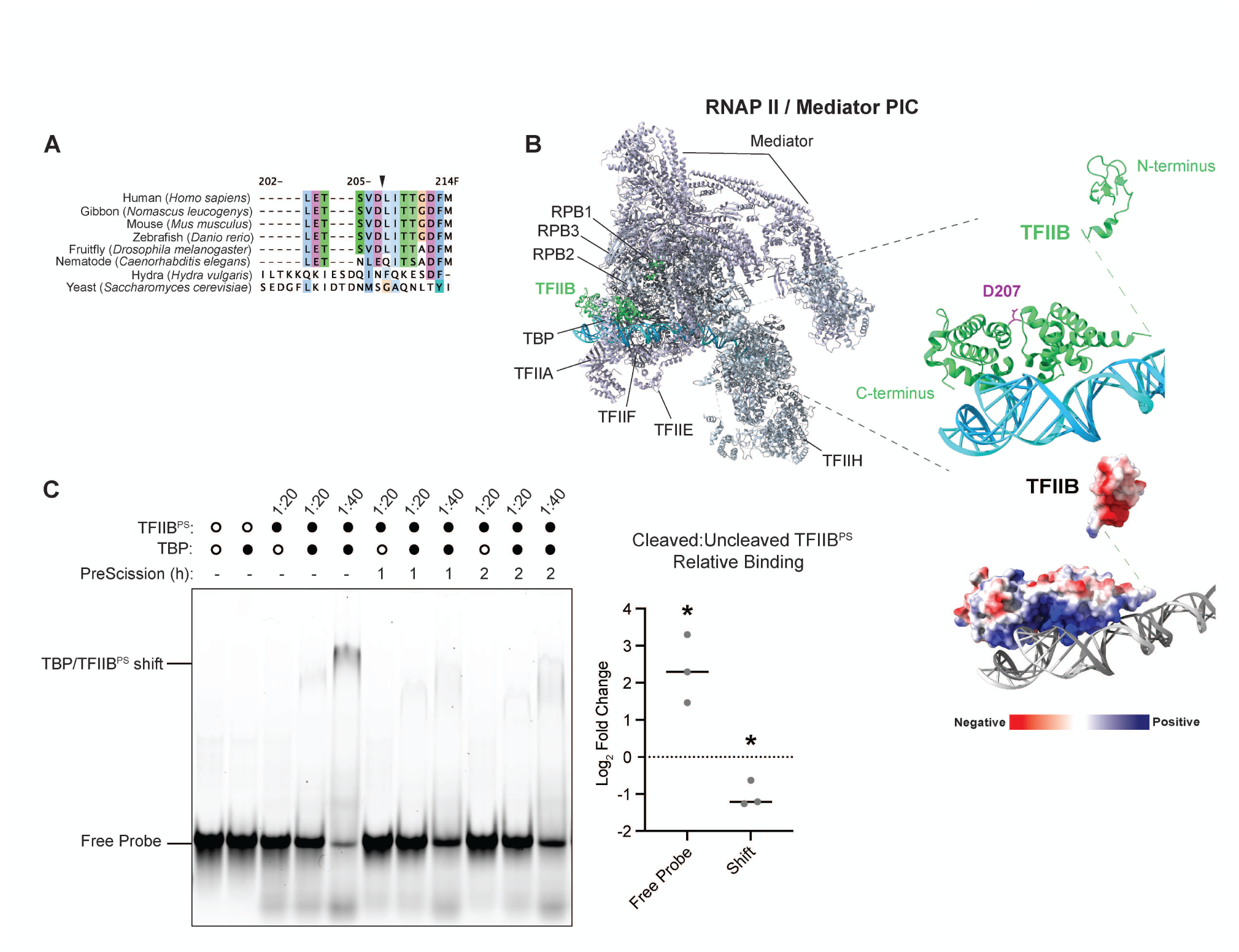
Cleavage after the conserved D207 residue impairs TFIIB promoter binding. **(A)** Multiple sequence alignment of the TFIIB protein near the D207 locus ranked by evolutionary time. The human sequence is used as a reference with dashes denoting missing areas of homology, and a black arrow indicates the precise location of the caspase cut site. Amino acids are colored with Clustal X scheme in Jalview to represent properties and conservation. **(B)** Structure of the RNAP II / Mediator pre-initiation complex from PDB:7NVR (Aibara et al. 2021) with TFIIB highlighted in green and promoter DNA in blue. Popout: D207 residue is highlighted in magenta between the TFIIB cyclin folds contacting DNA; shown beneath is the electrostatic overlay with increasing negative charge in red and increasing positive charge in blue. **(C)** EMSA of TFIIB with a PreScission site inserted after D207 (TFIIB^PS^) pre-incubated with PreScission protease for 1 h before (2 h total) or added with DNA for a 1 h binding reaction with Cy5-labelled AdML promoter DNA +/- TBP. The molar ratio of DNA:TFIIB^PS^ used is indicated above the lane. Graph on the right shows quantitation of the log_2_ fold change in free probe and shift for 40 molar TFIIB^PS^ EMSA reactions following 1 h PreScission incubation (cleaved TFIIB^PS^) relative to the uncleaved PreScission (-) control. \**p* < 0.05 in a one sample t and Wilcoxon test with a theoretical mean of 0.

### Failure to cleave TFIIB in cells upregulates transcription of a subset of genes during apoptosis

To define how caspase cleavage of TFIIB D207 regulates cellular gene expression during stress, we first used BCBL-1 cells stably overexpressing strep-tagged WT or uncleavable D207A versions of TFIIB. We performed RNA-seq on these cells after mock or TRAIL treatment to induce apoptosis, a condition in which mRNA synthesis is broadly decreased (Duncan-Lewis et al. 2021) **(Supplemental Fig. S3A-B)**. An exon-intron split analysis (Gaidatzis et al. 2015) was performed to quantify exonic and intronic read counts, and the resultant intronic counts were used to approximate differential expression of nascent transcripts **(Fig. 3A)**. This revealed 63 differentially expressed genes, nearly all of which (61) were upregulated in TFIIB D207-expressing cells. Although the number of changes is small, possibly due to higher overall levels of TFIIB in these overexpression lines, the trend is consistent with increased cellular availability of the cleavage-resistant form of TFIIB during stress.

**Figure 3.**
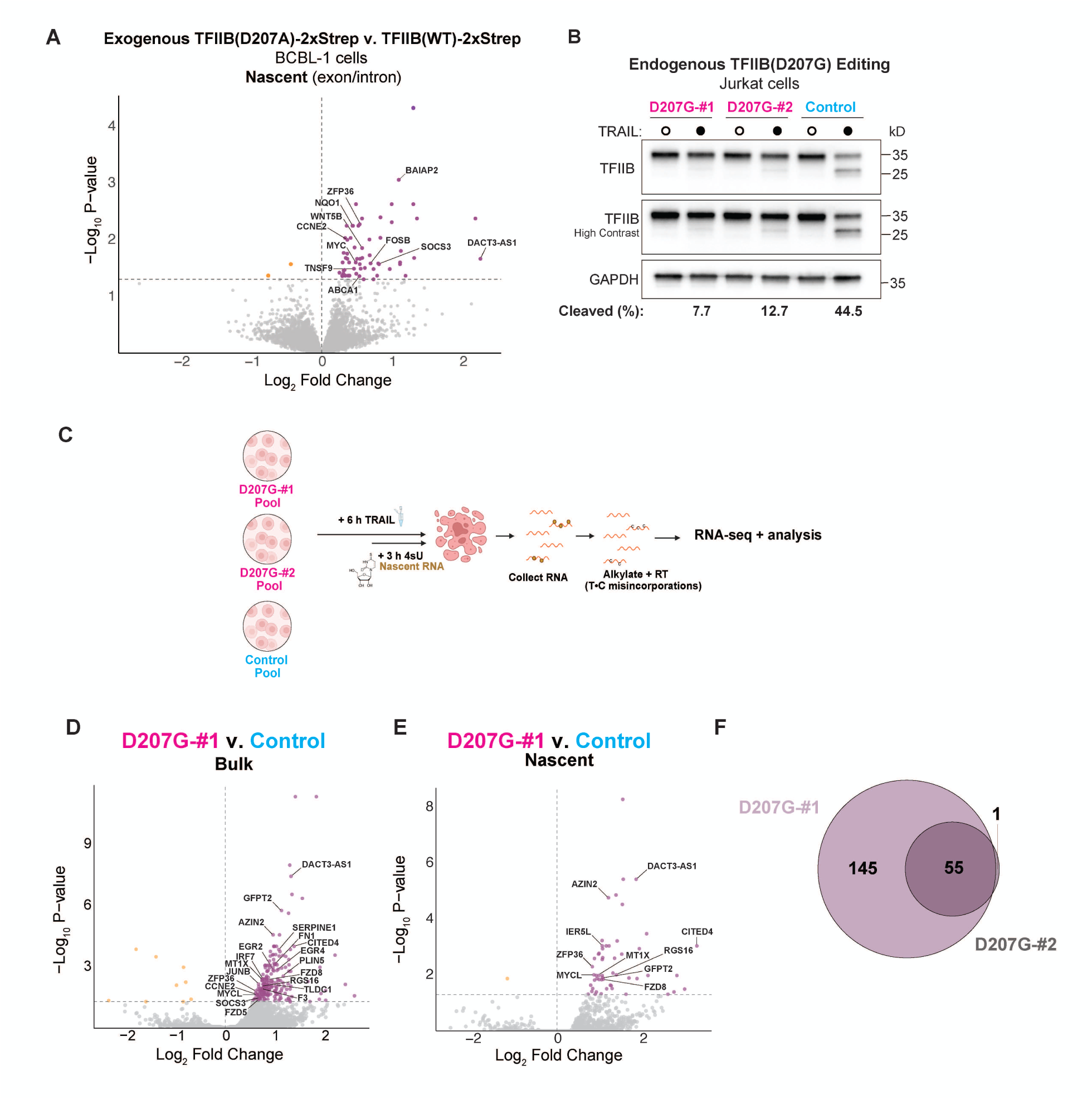
Expression of uncleavable TFIIB by transgene or endogenous editing in lymphocytes upregulates select genes during apoptosis. **(A)** Volcano plot showing nascent differentially expressed genes following an exon-intron split analysis from BCBL-1 cells overexpressing TFIIB(D207A)-2xStrep relative to cells overexpressing wild-type TFIIB-2xStrep after 7 h 660 ng/mL TRAIL treatment. Significantly downregulated genes are in yellow and significantly upregulated genes are in purple. **(B)** Western blots of control and D207G base edited Jurkat cell pools following treatment +/- 250 ng/mL TRAIL ligand for 6 h. Contrast is enhanced as indicated and GAPDH is a loading control. Percentage of total TFIIB protein that is cleaved is indicated for + TRAIL conditions. **(C)** Schematic for SLAM-seq (RT = reverse transcription). **(D-E)** Volcano plot showing bulk (*D*) or nascent (*E*) RNA differential gene expression between the TFIIB D207G-#1 guide and the control guide populations. Significantly downregulated genes are in yellow and significantly upregulated genes are in purple. TRAIL-specific gene upregulations: 200 bulk, 55 nascent. **(F)** Venn diagram overlap of genes upregulated in bulk RNA from D207G-#1 and D207G-#2 versus control guide populations. Genes differentially changed under mock conditions are excluded.

To avoid complications associated with overexpression, we next endogenously base edited TFIIB residue 207 to D207G in Jurkat T cells, a model cell line for studying apoptosis. Two different guides were used (D207G-#1 and D207G-#2); a guide targeting the AAVS1 safe locus served as a control (Lombardo et al. 2011; Li et al. 2019). All guides yielded efficient (≥ 85%) editing with minimal bystander effects, and the D207G mutation was stably maintained **(Supplemental Fig. S4A-D)**. Clonal lines with pristine edits were also derived from each guide; however, we primarily focused on the pooled lines, as clones are known to exhibit variation in gene expression due to drift (Meir et al. 2020). As expected, the D207G mutation substantially reduced cleavage of TFIIB in pooled populations and abolished cleavage in clonal lines treated with TRAIL relative to control **(Fig. 3B, Supplemental Fig. S5A).**

To rigorously quantify changes to both bulk and nascent RNA during apoptosis, each of the edited lines and matched controls was mock- or TRAIL-treated and subjected to SLAM-seq (thiol (SH)-Linked Alkylation for the Metabolic Sequencing of RNA) **(Fig. 3C)** (Jürges et al. 2018). The primary phenotype of TRAIL-treated D207G Jurkat cells was a significant increase in the expression of a subset of genes (∼50-200) in both bulk and nascent RNA relative to TRAIL-treated control cells, despite global transcriptional repression **(Fig. 3D-E; Supplemental Fig. S5B-C)**. Perhaps not surprisingly, there were a greater number of changes in the endogenously edited Jurkat cells compared to the BCBL-1 cells overexpressing tagged TFIIB. Notably, the quantity of upregulated genes scaled with the editing efficiency of the guide **(Fig. 3D-E** for the better edited D207G-#1 pool, **Supplemental Fig. S5D-E** for D207G-#2), and we observed nearly complete overlap in the set of TRAIL-specific genes that were significantly upregulated **(Fig. 3F)**. TFIIB D207G did not globally rescue TRAIL-induced transcriptional repression, which agrees with the fact that not all TFIIB protein was cleaved and indicates that other mechanisms of repression are also operational under these conditions (Duncan-Lewis et al. 2021). Nonetheless, a subset of genes is particularly sensitive to changes in TFIIB abundance in the context of TRAIL-treatment, and their RNA levels are reduced by stress-induced TFIIB cleavage.

### TFIIB downregulation by caspases specifically targets the expression of induced stimulus response genes

We next sought to understand what general characteristics define the TFIIB cleavage-sensitive loci identified above. We first noted that these genes are significantly shorter than all genes and their downregulated counterparts in both the Jurkat and BCBL-1 datasets, indicating that TFIIB stabilization disproportionally enhances expression of short transcripts **(Fig. 4A, Supplemental Fig. S6A)**. This is often a feature of stimulus-responsive genes, which tend to be associated with immune-related and/or proto-oncogenic functions (Tullai et al. 2007; Lopes et al. 2021). Some of the D207A/G-upregulated genes were the same across both the B and T cell lines (e.g, *ZFP36*, *AZIN2*, *PLIN5*, *CCNE2*, *SOCS* family genes). In many other cases the specific gene identities differed between Jurkat and BCBL-1 cells but nonetheless fell within similar categories. These include canonical immediate-early (IE) and delayed-early (DE) genes like *MYC, FOSB,* and *RGS1* in BCBL-1s **(Fig. 3A, Supplemental Fig. S3B)**, and *IER5L*, *GFPT2, JUNB, SERPINE1, EGR4* in Jurkat cells **(Fig. 3D-E, Supplemental Figs. S5D-E and S6B-C**). Other genes linked to cell proliferation and apoptosis were upregulated in aggregate in either B or T cells, such as *TNFSF9, BCL2, MT1X, TGFB1,* and Wnt signaling components like *WNT5B* and *FDZ* receptor genes. For full gene lists, see **Supplemental Table S1**. A reproducible association with specific pathways was evident upon Molecular Signatures Database (MSigDB) enrichment analysis. In line with the upregulated gene patterns above, both BCBL-1 and Jurkat gene sets were enriched in stress response and pro-survival pathways, particularly TNF-α signaling via NF-κB and inflammatory responses **(Fig. 4B, Supplemental Fig. S6B-E**).

**Figure 4.**
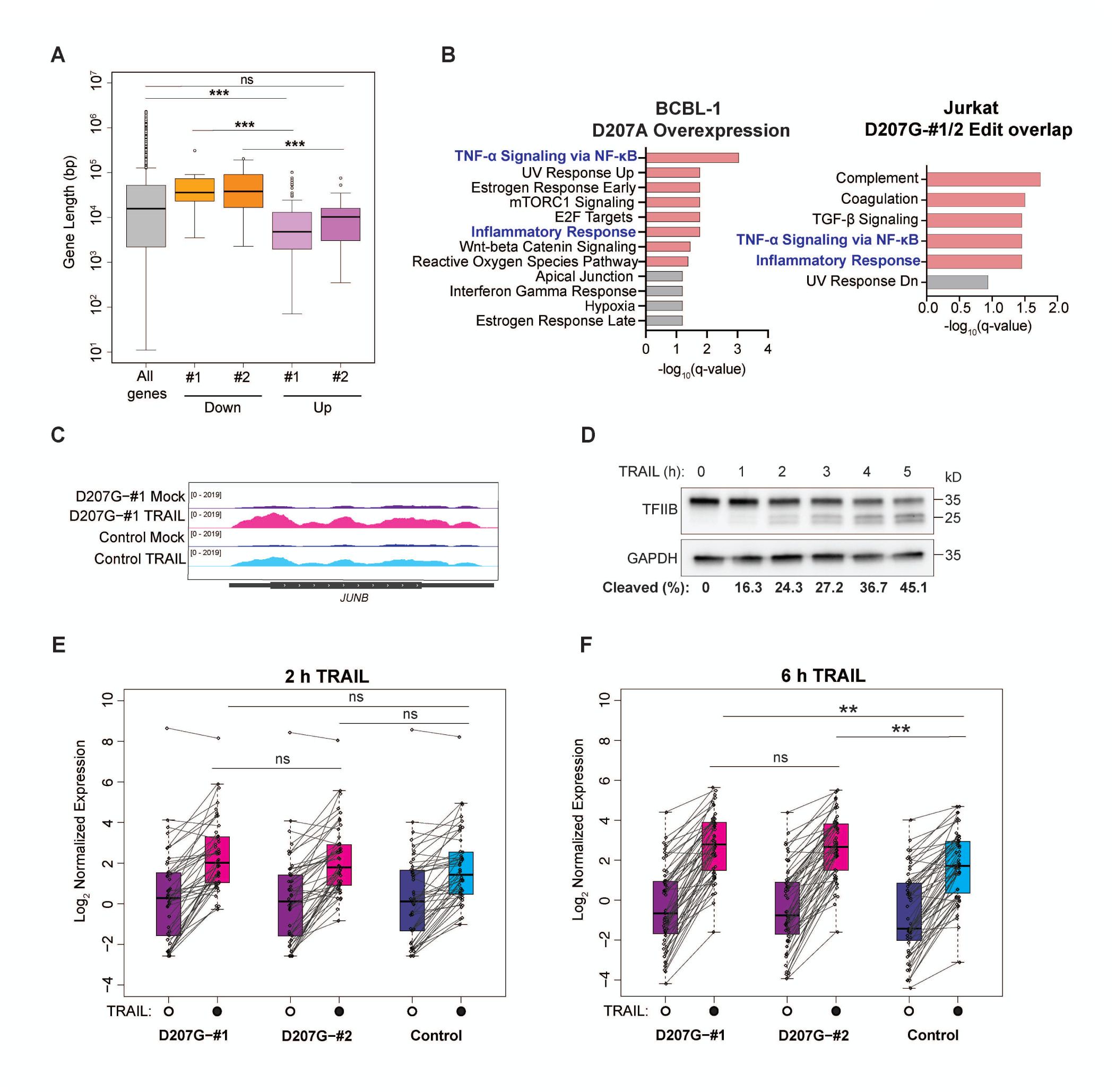
TFIIB cleavage tempers transcription from short, stimulus-responsive loci during apoptosis. **(A)** Boxplot comparing gene lengths for all genes detectable in the dataset (n = 23,806) and genes downregulated (yellow) or upregulated (purple) in a TRAIL-specific manner in bulk RNA from D207G-#1 or #2 guide cells. Significance values (Wilcoxon rank sum test with continuity correction) are relative to all genes; \*\*\**p* < 0.001; ns is nonsignificant. **(B)** Molecular Signatures Database Hallmark 2020 (MSigDB) enrichment from annotated genes (left) upregulated in nascent RNA in BCBL-1s overexpressing TFIIB(D207A)-2xStrep from Fig. 3A or (right) reproducibly upregulated in bulk RNA from D207G-#1 or #2 guide cells (overlap in Fig. 3E). Bars in pink are enrichments with a significant *q*-value while those in gray have significant *p*-values but nonsignificant *q*-values. **(C)** Bulk RNA coverage track of *JUNB* in D207G-#1 or control cells during mock or TRAIL-treatment. **(D)** Western blots of wild-type Jurkat cells following treatment +/-250 ng/mL TRAIL ligand for 0-5 h. Percentage of total TFIIB protein that is cleaved is indicated for each lane. **(E)** Jittered line and boxplot showing log2 normalized nascent RNA expression levels after 2 h TRAIL treatment for the 55 loci upregulated in bulk RNA from both D207G-#1 or #2 Jurkat pools (overlap in Fig. 3F). **(F)** As in *E*, but for log_2_ normalized bulk RNA expression after 6 h TRAIL treatment. Significance results comparing the three TRAIL conditions for *E-F* are shown for a one-way ANOVA with Tukey HSD; \*\**p <* 0.01, ns is nonsignificant. Comparisons amongst the three mock conditions are not shown but were nonsignificant.

The inducibility of these loci and pathways during cell perturbation appeared to be a key trait associated with TFIIB sensitivity. Indeed, the majority of the TFIIB cleavage-sensitive genes were apoptosis-induced in both WT and mutant cell lines but expressed at higher levels in D207G-edited cells (RNA-seq signal from a representative gene, *JUNB,* is shown in **Fig. 4C**). We thus hypothesized that TFIIB cleavage temporally restricts their transcriptional induction, perhaps to maintain their levels appropriately during stress. In this scenario, the genes should be similarly induced in WT and D207G cells shortly after the stress is applied, but their induction should be subsequently tempered by TFIIB cleavage in WT cells. To test this, we performed SLAM-seq at an early timepoint (2 h) during TRAIL treatment. This timepoint was chosen to enable sufficient time for induction of stress-response genes and because TFIIB cleavage then is relatively minimal, such that the amount of full-length TFIIB was similar in D207G and control cells **(Fig. 4D, Supplemental Fig. S7A-D)**. For stringency, we focused on the set of genes that was upregulated consistently between D207G-#1 and D207G-#2 Jurkat pools by 6 h of TRAIL exposure. Notably, at the 2 h timepoint, the majority of these genes were similarly induced and there was no significant difference in their overall expression between the D207G edited cells and the control cells **(Fig. 4E, Supplemental Fig. S7E)**. However, by 6 h of TRAIL treatment, the gene expression of this set was significantly enhanced in D207G-edited cells **(Fig. 4F, Supplemental Fig. S7F)**, in agreement with TFIIB cleavage limiting their synthesis as apoptosis progresses.

We also examined TFIIB promoter occupancy by chromatin immunoprecipitation (ChIP)-seq to determine whether its DNA-bound levels reduced after prolonged TRAIL treatment. A metagene analysis of TFIIB occupancy near the transcription start sites of TFIIB cleavage-sensitive loci showed a reduced peak by 4 h of TRAIL-treatment relative to 0 h control conditions (**Supplemental Fig. S8A-D)**. Some induced genes maintain poised RNAP II in unperturbed conditions (Ainbinder et al. 2002), which likely explains TFIIB occupancy at these loci in the 0 h control. Interestingly, discriminative MEME analysis of the regions upstream of loci that were nascently upregulated in D207A/G-expressing lymphocytes identified a GC-rich KLF and SP transcription factor family binding motif **(Supplemental Fig. S8E)**, as well as an enrichment in KLF4 and the negative elongation factor subunit NELFE targets **(Supplemental Fig. S8F)**. Since KLF4, SP1, and NELF are known to be involved in the regulation of stimulus-responsive genes, especially those mediated by NF-κB (Hirano et al. 1998; Ainbinder et al. 2002; Gilchrist et al. 2008; Fowler et al. 2011; Bahrami and Drabløs 2016; Ghaleb and Yang 2017), this points to possible underlying regulatory features that distinguish these sites.

Altogether, these data suggest that TFIIB cleavage plays a role in dampening the expression of short, responsive loci that are rapidly induced during stress and whose expression must be limited as the cell resolves stress or undergoes apoptosis (Bahrami and Drabløs 2016). Since many of these genes have mitogenic or pro-survival functions, we subsequently investigated how their overexpression dictates cellular outcomes during stress.

### Failure to cleave TFIIB causes aberrant proliferation of cells treated with etoposide and antimycin A

We hypothesized that failure to cleave TFIIB and subsequent upregulation of such pathways may cause aberrant cellular proliferation. We therefore performed a cell growth assay in which GFP-labelled control and mCherry-labelled D207G Jurkat cell pools were competed against each other in the presence of stress-inducing agents, including drugs with chemotherapeutic, anti-inflammatory, and neurological effects (**Fig. 5A**). The ratio of mCherry to GFP was calculated as a proxy for D207G cell fitness.

**Figure 5.**
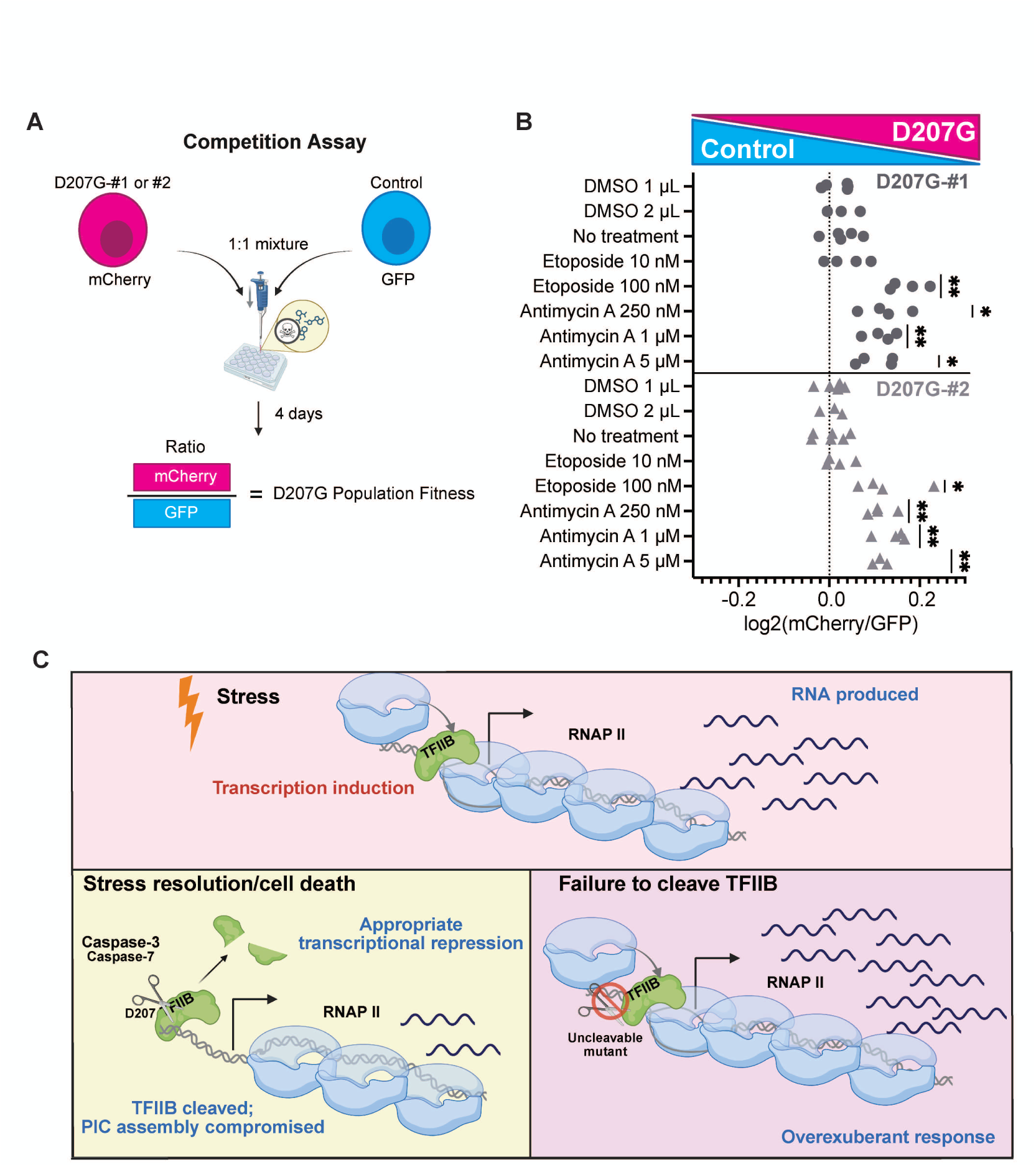
TFIIB D207G cells outcompete control cells in co-culture during chemical stress. **(A)** Schematic of the cell growth competition assay. Fluorescence measurements quantifying the relative percentage of mCherry and GFP cells were made via flow cytometry. Differences in plating and basal growth were accounted for by normalizing mCherry:GFP ratios to that of a DMSO control measured on Day 3. **(B)** Log2 normalized mCherry:GFP ratios for independent replicates of D207G-#1:control competitions (circles) and D207G-#2:control competitions (triangles). \**p* < 0.05, \*\**p* < 0.01; one sample t-test with theoretical mean of 0. **(C)** Schematic of model hypothesizing how TFIIB cleavage regulates stress-responsive genes.

Among the panel of chemical stressors tested, two candidates significantly altered the D207G:control ratio **(Supplemental Fig. S9A-B)** — the topoisomerase inhibitor etoposide, which is a chemotherapeutic agent used to treat various cancers, including those caused by KSHV, and the mitochondrial stressor antimycin A. The D207G cells outgrew the control cells in both cases and in a dose-dependent manner **(Fig. 5B, Supplemental Fig. S9C-D)**. This was not due to accelerated basal proliferation, as the D207G:control ratio in untreated and DMSO-treated populations did not increase over time. Thus, the inability to cleave TFIIB during stress can dispose cells to an abnormal state of proliferation.

## Discussion

Here, we uncover a mechanism by which cells dynamically alter protein abundance of the basal RNAP II general transcription factor TFIIB during a variety of stresses, including TRAIL treatment, DNA damage, translational stress, and KSHV lytic reactivation. Executioner caspases-3/7 cleave TFIIB at a highly conserved site located between its two cyclin folds, impairing its ability to interact with promoter DNA in vitro. This in turn tunes the cellular expression of a select set of stress-induced loci, presumably to temper transcription as required for stress resolution. Stressed cells unable to cleave TFIIB exhibit excessive transcription of these genes, a number of which are associated with pro-survival and mitogenic signaling. Functionally, we demonstrate that this can result in aberrant cellular proliferation during chemical stress **(Fig. 5C)**.

### Stress selectively reshapes transcription by targeting RNAP II protein machinery

Rapid responses to environmental stimuli are essential for viability. Repression of transcription may play a key role in shifting resources to activate a stress response or antiviral genes, as well as potentiate cellular shutdown when cell death is triggered. Caspase-mediated decay of TFIIB likely contributes to this by modulating the induction of a subset of stress-induced genes, many of which are short and have reported oncogenic activity. The sensitivity of only a specific subset of genes to TFIIB cleavage likely reflects the fact that (1) TFIIB loss by cleavage is incomplete and some residual full-length protein remains and (2) acute stress disrupts RNA synthesis through multiple pathways. For example, during genotoxic stress, the RNAP II catalytic subunit RPB1 is degraded by caspase- and ubiquitin-mediated pathways (Lu et al. 2002; Wilson et al. 2013; Caron et al. 2019; Tufegdžić Vidaković et al. 2020).

As previously noted for RPB1-sensitive genes, short and stimulus-induced genes may be less susceptible to the generalized transcriptional shutdown that occurs during stress because of a higher probability that initiating polymerases can successfully complete their transcription (Tufegdžić Vidaković et al. 2020). For these genes, impeding initiation by reducing available RNAP II machinery like TFIIB should be an effective means of controlling their expression. Whether this primarily occurs through diminished TFIIB abundance or if TFIIB cleavage can also destabilize assembled PICs remains an open question.

Our data support the hypothesis that the kinetics of TFIIB cleavage temporally influence transcriptomic shifts. While cells with cleavage-resistant TFIIB (D207G) resembled WT gene expression profiles after 2 h TRAIL treatment, they maintained higher expression levels later (at 6 h), coinciding with increased levels of WT protein cleavage. Curiously, the TSVD caspase-3/7 recognition motif within TFIIB does not adhere to the optimal DEVD recognition motif, nor does it contain the canonical negatively charged residue in the P4 position (Julien and Wells 2017). The non-canonical TS residues in the respective P4 and P3 locations may attenuate caspase recognition efficiency (Julien and Wells 2017) and potentially delay cleavage of TFIIB relative to other substrates in a controlled manner. Cells with less potent cell death pathways may consequently retain more TFIIB, leveraging its abundance as a rheostat to tune transcription.

### Physiological context of TFIIB regulation

B and T lymphocytes exhibit high rates of turnover (Young et al. 1995) and are strictly controlled by cell death at various stages, including during maturation and in the post-immune response “contraction” phase (Zhang et al. 2005; Shah et al. 2021). These cells exist under a fine balance of cell death and survival pathways (Thome and Tschopp 2001; Sen 2006), the dysregulation of which can lead to lymphoproliferative disorders (Justiz Vaillant and Stang 2025). Our finding that TFIIB(D207G)-edited cells have a competitive growth advantage during mitochondrial stress and DNA damage underscores the importance of this mechanism of TFIIB regulation. More broadly, TFIIB is a common essential gene for most cancer cell lines (Arafeh et al. 2025; DepMap 2025), and pan-cancer analysis suggests TFIIB-dependent metabolic genes are often dysregulated (Rosario et al. 2018). Of the ∼13 mutations reported throughout TFIIB by DepMap, one (E203) is proximal to the cleavage site and altered in some cancer lines, which could be examined in future work. Likewise, caspases (including caspase-3 and 7) are mutated or differentially regulated in some cancer tissues; as mediators of inflammation and cell death, modulating caspase activity is an area of interest for oncology (Ghavami et al. 2009).

It is worth noting that not all caspase activity results in apoptosis; processes such as development and differentiation also depend on caspases in nonlethal contexts (Nakajima and Kuranaga 2017). Thus, caspase-mediated regulation of TFIIB could dictate a range of cellular processes. This includes virus-host interactions, where control of TFIIB, and more broadly RNAPII, is an important component (O’Brien and Ansari 2021; Gulyas and Glaunsinger 2022). Because nuclear-replicating DNA viruses like KSHV must hijack cellular transcription for their own gene expression, how they contend with TFIIB cleavage and its potential role in restricting viral transcription are exciting questions for the future.

## Materials and Methods

Methods for some supplemental data are found in **Supplemental Materials and Methods**.

### Cloning

The human TFIIB sequence (synthesized by Twist Bioscience) for cloned constructs was modified to preserve the amino acid sequence but be resistant to Dharmacon OnTarget and Accell human *GTF2B* siRNAs. For strep-tagged constructs, TFIIB was Infusion (Takara Bio) cloned into XhoI site of pcDNA4/TO-2xStrep (pCDNA4/TO Invitrogen vector modified to include a C-terminal 2xStrep tag). TFIIB-2xStrep was Infusion cloned into the Sfi1 site of a sleeping beauty plasmid pSBbi-GP (Addgene #60511, gift from Eric Kowarz) derivative lacking EGFP to make the TFIIB-2xStrep expression vector (Addgene #213919). Derivative point mutant plasmids D207A, S65A, S65E, and R66E (Addgene #213920-2, 252053) were made by inverse PCR as described previously (Silva et al. 2023). The siRNA resistant 6xHis-TFIIB-V5 (Addgene #242714) construct was synthesized by Twist, amplified, and InFusion subcloned with a PCR-amplified sleeping beauty plasmid pSBtet-BB backbone (Addgene #60505, gift from Eric Kowarz). Sleeping beauty vectors and the sleeping beauty SB100X transposase plasmid were provided by Nicholas Ingolia (UC Berkeley); the sleeping beauty system was described previously (Kowarz et al. 2015).

All plasmids were validated by full-plasmid sequencing and deposited on Addgene. Primer and DNA sequences are available in **Supplemental Table S2**.

### Cell line generation, maintenance, and reactivation

All cell lines were maintained at 37°C and 5% CO_2_ in a humidity-controlled incubator, and cell density was kept at 0.5-2 × 10^6^ cells/mL. TRExRTA BCBL-1 cells (generously provided by the lab of Jae Jung) (Nakamura et al. 2003) were maintained in RPMI 1640 media (Gibco) supplemented with 10% Tet-free fetal bovine serum (FBS; via UC Berkeley Cell Culture Facility), 1X GlutaMAX (Gibco), and 10 μg/mL hygromycin B (Invitrogen). Unless otherwise noted, cell concentration was ∼1 × 10^6^ cells/mL when reactivated with 1 μg/mL doxycycline or drug-treated. Jurkat cells (obtained from the UC Berkeley Cell Culture facility) were maintained in RPMI 1640 media (Gibco) with 1X GlutaMAX (Gibco), 10 % Tet-free FBS (via UC Berkeley Cell Culture Facility), 1 mM sodium pyruvate, and 1X Pen-Strep (Gicbo).

BCBL-1 derivatives (aside from the R66E mutant, which was transiently nucleofected with 0.5 μg vector DNA into 3 × 10^6^ cells for 18 h) were generated by nucleofection with the Neon Transfection System (Invitrogen) according to the manufacturer’s instructions with the following parameters: 5 × 10^6^ low-passage cells were pulsed (1350 V, 1 × 40 ms pulse-length) with 2 μg vector DNA and 0.1 μg transposase plasmid in a 100 μL tip using resuspension buffer T. Cells were selected with 1 μg/mL puromycin 24 h post-nucleofection.

### Jurkat base editing and labeling

Cells were base edited by nucleofecting (Neon Transfection System per manufacturer’s instructions; 1700 V, 1 × 20 ms pulse-length, 10 μL tip) 1× 10^6^ low passage Jurkat cells with 1.63 µg A•G base editor RNA (Gaudelli et al. 2017) and 1 μg sgRNA (IDT) containing target sequences (**Supplemental Table S2**) predicted by the UCSC Genome Browser (Perez et al. 2025). The AAVS1 safe sequence was previously published (Li et al. 2019). The in vitro-transcribed ABE8e base editor RNA from Addgene #138489 (gift of David Liu) was generously provided by Fyodor Urnov (UC Berkeley).

mCherry (pLVX-mCherry-P2A-Blasticidin) and GFP (pLVX-GFP-P2A-Blasticidin) (Haakonsen et al. 2024) labelling plasmids were generously provided by Michael Rapé (UC Berkeley). Labelling of base-edited pooled populations was performed via 2^nd^ generation lentiviral transduction: 1 × 10^6^ Lenti-X 293T cells (Clonetech/Takara via the UC Berkeley Tissue Culture Facility) maintained in DMEM (Gibco) with 10% FBS were transfected (manufacturer’s protocol) using Mirus *Trans*IT-293 Transfection Reagent with 250 ng pCMV-VSV-G (Addgene #8454, gift of Bob Weinberg), 1250 ng psPAX2 (Addgene #12260; gift of Didier Trono), and 1250 ng lentivector. Lentivirus was harvested at 48 h, 0.45 μm filtered, and 3 mL of viral product was mixed with 5 × 10^6^ base edited Jurkat cells in 2 mL of media (D207G guide populations received mCherry; AAVS1 control received GFP), and 10 μg/mL polybrene (Sigma). Cells were spinfected at 800 × g for 2 h at 32°C, resuspended in 5 mL fresh media, recovered for 48 h, and selected with 5 μg/mL blasticidin HCl.

### siRNA knockdowns

3 × 10^6^ BCBL-1 cells were nucleofected (1350 V, 1 × 40 ms pulse-length) in 100 μL of resuspension buffer T (Invitrogen) with the Neon Transfection System (Invitrogen) according to the manufacturer’s instructions. 3 μM of either Dharmacon human *Casp3* and/or *Casp7* Accell SmartPool siRNA and 3 μM Dharmacon human OnTarget *Casp3* and/or *Casp7* SmartPool were used. Non-targeting Accell and OnTarget siRNA (siNTC) was used as a control.

### Protein purification and in vitro assays

A pET-28a(+) expression vector containing 6xHis-SUMO-TFIIB^PS^ (in which a LEVLFQ/GP PreScission protease recognition site was introduced directly after the TFIIB D207 residue) was synthesized by Twist Bioscience (Addgene #252054). After transformation, NiCo21(DE3) Competent *E.coli* (NEB) were grown in 1 L Terrific Broth (ThermoFisher) at 37°C until OD_600_ ∼1.1 and induced with 0.2 mM IPTG overnight at 18°C. Conditions for purification are in Buffer A (25 mM Tris-HCl pH 7.5, 10% glycerol, 5 mM 2-mercaptoethanol) at indicated salt concentrations unless otherwise noted. The cell pellet was lysed by sonication (qSonica q700 macrotip; 11 min, 50 A, 15 s on/45 s off) in 60 mL of Buffer A (200 mM NaCl, Halt Protease and Phosphatase Inhibitor Cocktail (ThermoFisher)). The lysate was clarified by centrifugation (25,000 × g, 45 min, 4°C) and the supernatant applied to a 2 mL Ni-NTA agarose bed (ThermoFisher) column equilibrated with 10 CV lysis buffer. The column was washed with 10 CV Buffer A (1M NaCl) + 10 mM Imidazole-HCl pH 7.5, then with 10 CV Buffer A (150 mM NaCl) + 20 mM Imidazole-HCl pH 7.5. Sample was eluted in 1 mL increments with Buffer A (150 mM NaCl) + 200 mM imidazole-HCl pH 7.5. TFIIB-containing fractions were pooled with > 1:50 molar TEV protease:protein (Berkeley QB3 Macrolab) and dialyzed overnight with a 10,000 MWCO Slide-a-Lyzer dialysis cassette (ThermoFisher) in Buffer A (pH 7.8, 100 mM NaCl). TFIIB^PS^ was purified by Ni-NTA agarose gravity column, concentrated to 3 mL in an Amicon Ultra-15 10,000 MWCO Centrifugal Concentrator, then passed over a 5 mL HiTrap Q HP column (Cytiva ÄKTA Pure chromatography system running Unicorn 7.0) in the presence of 5 mM DTT to remove residual impurities. The column flowthrough containing TFIIB^PS^ was reconcentrated and loaded onto a HiLoad Superdex S200 pg gel filtration column (GE Healthcare Life Science) and eluted in storage buffer (20 mM Tris-HCl, pH 8.0, 100 mM KCl, 20% glycerol, 5 mM DTT, and 0.2 mM EDTA). Pure TFIIB^PS^ fractions were concentrated to 0.5 mg/mL and stored at -80°C. Wild-type human TFIIB and TBP were purchased in the same storage buffer and similar concentrations from ProteinOne.

For the EMSAs, TFIIB or TFIIB^PS^ was incubated at 30°C for the indicated times and molar ratios with or without 3.5 × 10^-3^ U PreScission Protease (Genscript). This was performed in 10 μL containing a total of 4 μL storage buffer and 10 μL EMSA buffer (40 mM Tris-HCl pH 8.0, 4 mM MgCl_2_, 5 mM (NH_4_)_2_SO_4_, 4% Ficoll, 2% PEG 8000 (w/v), 55 mM KCl, 0.1 mM EDTA, 1 mM DTT) to account for differences in required protein volume. After PreScission incubation, an additional 10 μL of EMSA buffer containing TBP and dsDNA probe were added to a final concentration of 200 nM TBP and 50 nM probe and incubated for another 1 h. A duplexed, 5’ sense Cy5-labelled modeled on the AdML promoter sequence (IDT; Sense: 5’-AAGGGGGGCTATAAAAGGGGGTGGGGGCGCGTTCGTCCTCACTCTCTTCCCCTCC ATACC-3’) served as a probe (Aso et al. 1994; Glossop 2004). To reduce activity from a minor nuclease contaminant in purified TFIIB^PS^, 35 nM of random ssDNA (**Supplemental Table S2**) was also included in all reactions. Reactions were mixed with 0.5 μL 6x Orange Loading Dye (ThermoFisher) and run on a 5% Criterion TBE polyacrylamide gel (Bio-Rad) in 0.5X TBE at 100 V (4°C). Gels were scanned at 50 μm on an Amersham Typhoon Imager (Cytiva) and analyzed with Bio-Rad Image Lab v. 6.0.1 to obtain the adjusted band volume (intensity) used for shift and free probe quantitation. The log_2_FC was determined from the band ratio in the PreScission (+) to PreScission (-) control at the same molar ratio of protein.

### Western blotting

Cells were lysed in 1% SDS RIPA buffer (1% SDS, 1% NP40, 150 mM NaCl, 50 mM Tris-HCl pH 8, 0.5% sodium deoxycholate by mass) with Halt Protease and Phosphatase Inhibitor Cocktail (ThermoFisher), followed by sonication (qSonica cuphorn; 1 min, 100 A, 3 s on/17 s off), 1% Benzonase (Millipore) treatment, and centrifugation (21,000 x g, 10 min, 4°C). Diluted samples were quantified by Pierce 660 assay with 0.8% Triton-X. 10-30 μg lysate was denatured in Laemmli Buffer (Bio-Rad), resolved by SDS-Page and transferred to PVDF membrane with a Trans-Blot Turbo apparatus (Bio-Rad, Mixed Molecular Weight protocol). Western blotting was performed with Tris-buffered saline and 0.2% Tween-20 and the following primary and secondary antibodies: TFIIB (clone 2F6A3H4, Cell Signaling- 4169, 1:1000), GAPDH (anti-mouse, clone 6C5, Thermofisher-AM4300, 1:5000 or anti-rabbit, clone 14C10, Cell Signaling-2118, 1:1000), Vinculin (Abcam- ab91459, 1:1000), Caspase-3 (Cell Signaling-9662, 1:1000), Caspase-7 (Cell Signaling-9492, 1:1000), V5 (clone D3H8Q, Cell Signaling-13202), Goat Anti-Mouse IgG(H+L) or Goat Anti-Rabbit IgG(H+L) Human ads-HRP (Southern Biotech, 1:5000), Strep-Tag II Antibody HRP Conjugate (Sigma Aldrich-71591-3, 1:2000). Blots were developed with Clarity Western ECL Substrate (Bio-Rad) or Radiance Plus Femtogram-sensitivity HRP Substrate (Azure Biosystems). Blots were imaged on a Chemidoc MP imager (Bio-Rad) and analyzed/prepared for publication using Bio-Rad Image Lab v. 6.0.1 with the autoscale feature to maintain consistency unless otherwise noted; blot quantitation was performed where applicable with the adjusted band volume (intensity) output. Blots shown are representative of a minimum of 3 biological replicates at the same conditions and/or timepoints.

### RNA-sequencing

RNA-seq experiments were conducted in triplicate as follows with the noted specifications. Cells were washed and pelleted, then total RNA was collected in 1 mL TRIzol (Invitrogen) and isolated by 1 × 200 uL chloroform extraction as outlined previously (Toni et al. 2018), including precipitation in 500 uL isopropanol with 1% glycogen and 10% 3 M sodium acetate. RNA concentration was quantified by Qubit 3 Fluorometer with High Sensitivity RNA Kit (Invitrogen). Prior to library prep, ERCC RNA Spike-in mix (Invitrogen) was added to all input samples per the manufacturer’s instructions. TFIIB-2xStrep/TFIIB(D207A)-2xStrep libraries were prepped by the KAPA RNA Hyperprep Kit with RiboErase (Roche) using 1,000 ng input RNA, with fragmentation at 94°C for 6 min, ligation with KAPA Unique Dual-Indexed adaptors (Roche), and amplification for 7 cycles. Library quality was confirmed on an AATI (now Agilent) Fragment Analyzer. Library molarity was measured via quantitative PCR with the KAPA Library Quantification Kit (Roche KK4824) on a Bio-Rad CFX Connect thermal cycler. Libraries were then pooled by molarity and sequenced, targeting at least 25M reads per sample. TFIIB-2xStrep/TFIIB(D207A)-2xStrep libraries were sequenced on an Illumina NovaSeq X 10B with 150 bp paired-end reads. Fastq files were generated and demultiplexed using Illumina bcl_convert and default settings, on a server running CentOS Linux 7.

Analysis was modified from the UC Davis Bioinformatics Core RNA-seq workflow. Sequencing quality was assessed with MultiQC, and reads were preprocessed with HTStream v. 1.3.0, including deduplication. Genome indices were prepared using STAR v. 2.7.1a (Dobin et al. 2013). The human GRCh38.p13 genome assembly (ensembl.org) was indexed with Gencode v43 annotations. Preprocessed reads were aligned with STAR, and count files were generated for transcripts. Nascent cellular RNA counts were estimated by the eisaR package as described (Gaidatzis et al. 2015). Library size normalization (TMM method) was used since bulk RNA of the same treatment showed minimal variance. Differential gene expression analysis for bulk and nascent RNA was conducted in RStudio v. 2024.12.1+563 with the Bioconductor package using edgeR with limma/voom developed and described by Law et al. (Law et al. 2018). Bulk RNA was normalized to ERCC Spike-In and multiple testing was compensated for using the Benjamini-Hochberg correction. Gene annotations are from the BioMart Ensembl Genes 109 database. Plots and statistics were generated with R (v. 4.3.2) ggplot2 and VennDiagram. Enrichment analyses are from Molecular Signatures Database Hallmark 2020 and Encode and ChEA Consensus TFs from ChIP-X via EnrichR (Liberzon et al. 2011; Chen et al. 2013; Kuleshov et al. 2016; Xie et al. 2021).

#### SLAM-sequencing

SLAM-seq was performed as above but with the following modifications. After 3 h treatment of SUPERKILLER TRAIL (250 ng/mL; Enzo) or mock diluent, 300 μM of 4-thiouridine (4sU) or PBS control was added to 3 × 10^6^ cells. Cells were harvested 3 h post-4sU for RNA or western blot. The 2 h SLAM-sequencing was performed identically, but both 4sU and TRAIL were added simultaneously for 2 h total before harvesting cells. RNA extraction (2 × 200 uL chloroform extraction) and alkylation were performed using the Lexogen SLAM-seq Anabolic Kinetics Kit according to the manufacturer’s instructions with 2 μL GlycoBlue (Invitrogen) as a co-precipitant. RNA integrity was verified by TapeStation (Agilent) and quantified using Qubit Broad Range Sensitivity RNA reagents. ERCC RNA Spike-in mix (Invitrogen) was added to inputs per the manufacturer’s instructions along with 5 ng of *in vitro* transcribed mixture of both unlabeled and 4sU-labeled spike-ins (synthesized as described previously (Schwalb et al. 2016; Gressel et al. 2019)).

Inputs for pooled (1000 ng) and clonal (700 ng) line sequencing were alkylated, then samples were DNase-treated and repurified with RNA Clean and Concentrator kits (Zymo). Quality and concentration were reassessed by TapeStation and Qubit (∼500 ng final input). Library preparation was performed with fragmentation at 94°C for 6 minutes. Pooled line samples were ligated with 3 μM adaptor and amplified for 9 PCR cycles; clonal samples were ligated with 1.5 μM and amplified for 10 cycles. Samples were sequenced on a Novaseq X Plus 25B with 150 bp paired-end reads. Incorporation of 4sU was validated by visualization on IGV (Robinson et al. 2011). Nascent RNA counts were estimated using GRAND-SLAM with default parameters (Jürges et al. 2018). Total count values were multiplied by the MAP statistic (nascent-to-total ratio) to yield approximate nascent counts; inestimable values were considered 0. Nascent and bulk RNA counts were analyzed using edgeR and limma/voom with ERCC Spike-in normalization from bulk RNA. RNA coverage plots were generated using deepTools (Ramírez et al. 2016) bamCoverage with spike-in normalized --scaleFactor inputs. BigWig files for biological triplicates were averaged with bigwigAverage (--binSize 1) and visualized in IGV.

Raw data and counts tables for RNA-seq experiments are available on the GEO database at accession numbers GSE250393, GSE302704, GSE302707, and GSE16831.

### Competition assays

1.5 × 10^5^ cells each of mCherry-labelled D207G pool and GFP-labelled control pool were co-cultured in 1 mL media with the indicated times and drug concentrations (dissolved in DMSO). Population fluorescence was assessed by flow cytometry on a BD Bioscience LSR Fortessa. 10,000 events/sample were collected, and replicates represent independent competitions on separate days. All data were analyzed using FlowJo software (BD Bioscience) with gating for live cells and PE-Texas-Red or FITC positivity. Normalization is indicated in figure legends.

### Data analysis and data visualization

Data quantifications, statistical analyses, and/or visualization for competition assays, EMSAs, and enrichment were conducted in GraphPad Prism v. 9-10.10; statistical analyses for gene length and RNA-seq expression changes were conducted in R (v. 4.3.2). The Multiple Sequence Alignment (MSA) was generated from TFIIB protein sequences obtained through UniProt (uniprot.org) and analyzed in JalView v. 2.11.4.1 (Waterhouse et al. 2009; Troshin et al. 2011; Troshin et al. 2018) with a human reference and Clustal X coloring/annotation. Protein structures were visualized in UCSF ChimeraX v. 1.9 (Pettersen et al. 2021). All schematics were created in BioRender (Gulyas, L. (2026) https://BioRender.com/sh8ifqb).

## Acknowledgements

We thank the reviewers and all members of the Glaunsinger and Coscoy labs (UC Berkeley) for their helpful advice and feedback, especially Dr. Chad Stein, Xiaowen Mao, and Sam Rider. We sincerely appreciate the training and support for protein purification and in vitro experimentation from Michael Palo (Stanford University) and Dr. Aaron Mendez (Tufts University). We are also grateful to the following UC Berkeley colleagues: Dr. Fyodor Urnov and Jessica Nguy (Urnov lab) for providing expertise in base editing and A•G base editor RNA, Dr. Michael Rapé and Dr. Michael Heider (Rapé lab) for providing helpful advice and reagents for the drug screen, and Dr. Annan Cook for useful discussions. We thank Dr. Florian Erhard (University of Regensberg) for support using GRAND-SLAM. Finally, we thank the UC Berkeley IGI NGS facility, UCSF ChimeraX (funded by NIH R01-GM12932, Office of Cyber Infrastructure and Computational Biology, NIAID), UC Davis Bioinformatics Core (RRID:SCR_023887) and the UC Berkeley QB3 Genomic Facilities (RRID:SCR_022170) and Macrolab for their services. This research was supported by NIH grants AI122528 and CA136367 to B.G., who is also an investigator of the Howard Hughes Medical Institute.

## Author contributions

Conceptualization, L.G. and B.A.G.; methodology, L.G. and B.A.G.; experimentation, L.G., A.L., S.B.S; writing, L.G. and B.A.G.; formal analysis, L.G., A.L, S.B.S., and B.A.G.; project administration, L.G. and B.A.G; supervision and funding acquisition, B.A.G.

