## Supplemental Figures and Methods for "Cleavage of the RNA polymerase II general transcription factor TFIIB tunes transcription during stress"

#### Supplemental Materials List

1. Supplemental Figures S1, S2, S3, S4, S5, S6, S7, S8, S9
2. Supplemental Materials and Methods and References (for Supplement-specific experiments)
3. Supplemental Table S1 (Upregulated gene lists): Lists of genes significantly upregulated in cells expressing uncleavable TFIIB during TRAIL treatment relative to wild-type TFIIB. Each spreadsheet tab has the list from the indicated comparison. (.xlsx)
4. Supplemental Table S2 (DNA sequences, primers): DNA sequences, primers, and sgRNA sequences used in this study. (.xlsx)

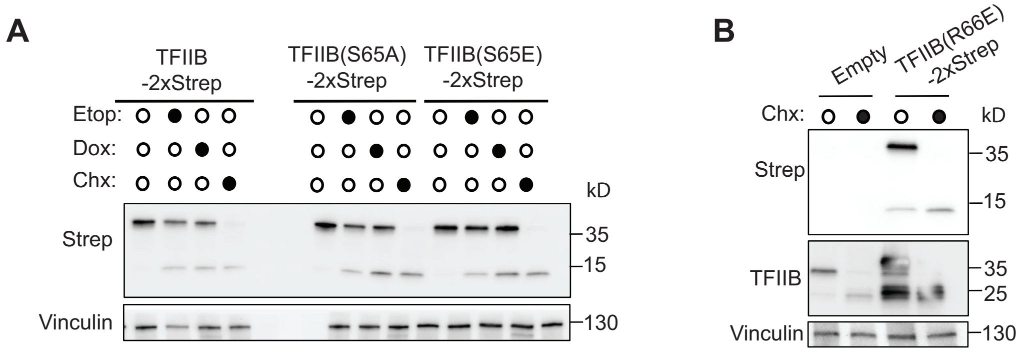


**Supplemental Figure S1. (A)** TRExRTA BCBL-1 cells expressing TFIIB-2xStrep, TFIIB(S65A)-2xStrep, or TFIIB(S65E)-2xStrep were mock treated, exposed to 25 μM etoposide (Etop) for 24 h, 2 μg/mL doxycycline (Dox) for 24 h, or 100 μg/mL cycloheximide (Chx) for 6 h, and analyzed by western blotting. Anti-TFIIB antibody recognizes both endogenous and strep-tagged constructs. Vinculin is a loading control. **(B)** Western blot of BCBL-1 cells transiently nucleofected with empty or TFIIB(R66E)-2xStrep expression vector for 18 h and treated with 100 μg/mL Chx for 6 h.

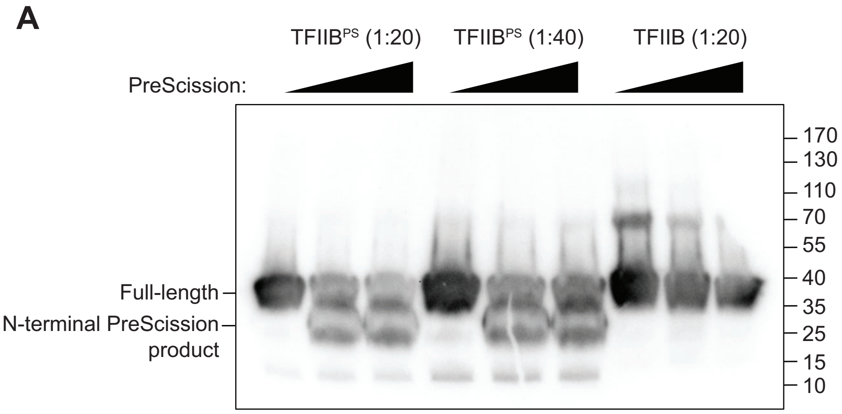


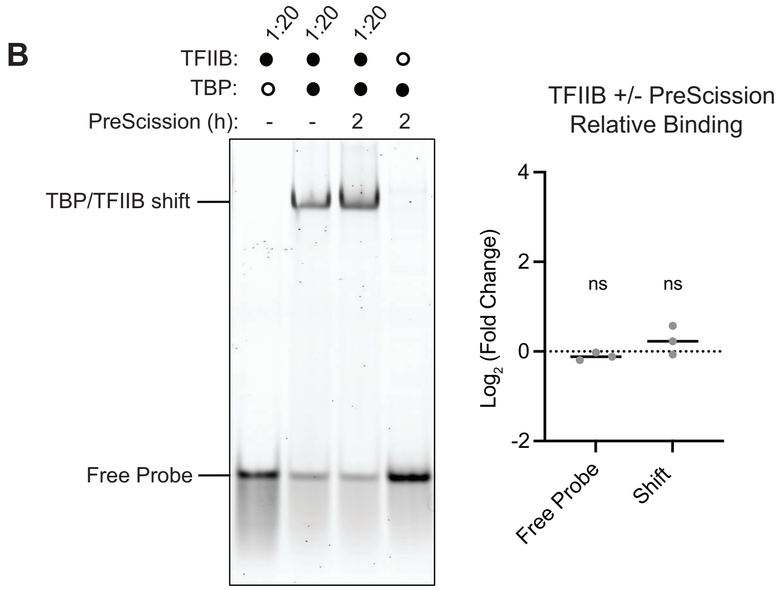


**Supplemental Figure S2. (A)** TFIIB$^{PS}$ or wild-type TFIIB at the molarity (20 or 40 molar excess relative to DNA) used for subsequent EMSA assays was incubated with 0, 3.5 × $10^{-3}$ U, or 7 × $10^{-3}$ U PreScission protease for 1.5 h and analyzed by western blot. Anti-TFIIB antibody detects the N-terminal portion of the protein, and the full-length and N-terminal cleavage product are indicated. **(B)** Wild-type TFIIB lacking the engineered PreScission site was incubated for 1 h with AdML promoter DNA and TBP +/- 1 h pre-incubation with PreScission protease (2 h total PreScission) and analyzed by EMSA. Graph on the right shows quantitation of the $log_2$ fold change in free probe and shift is for the reaction with PreScission relative to a PreScission (-) control. A 1:20 molar ratio of DNA:TFIIB was assessed as it produced a shift comparable to that observed for the 1:40 DNA:TFIIB$^{PS}$ quantified in main **Fig. 2C**. ns = nonsignificant in a one sample t and Wilcoxon test with a theoretical mean of 0.

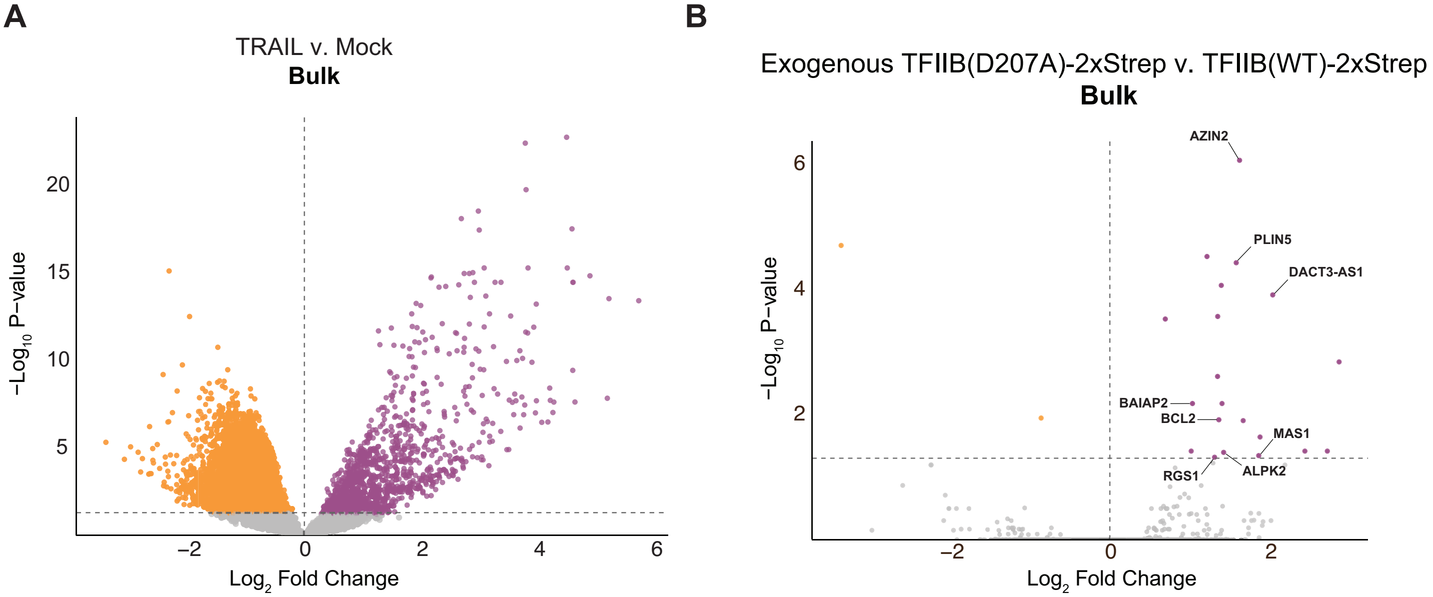


**Supplemental Figure S3. (A)** Volcano plot showing differentially expressed genes in bulk RNA during mock or 660 ng/mL TRAIL treatment of BCBL-1 cells expressing wild-type (WT) TFIIB-2xStrep. Significantly downregulated genes are in yellow and significantly upregulated genes are in purple. **(B)** As in *A* but for TRAIL-treated cells expressing the D207A mutant protein versus WT.

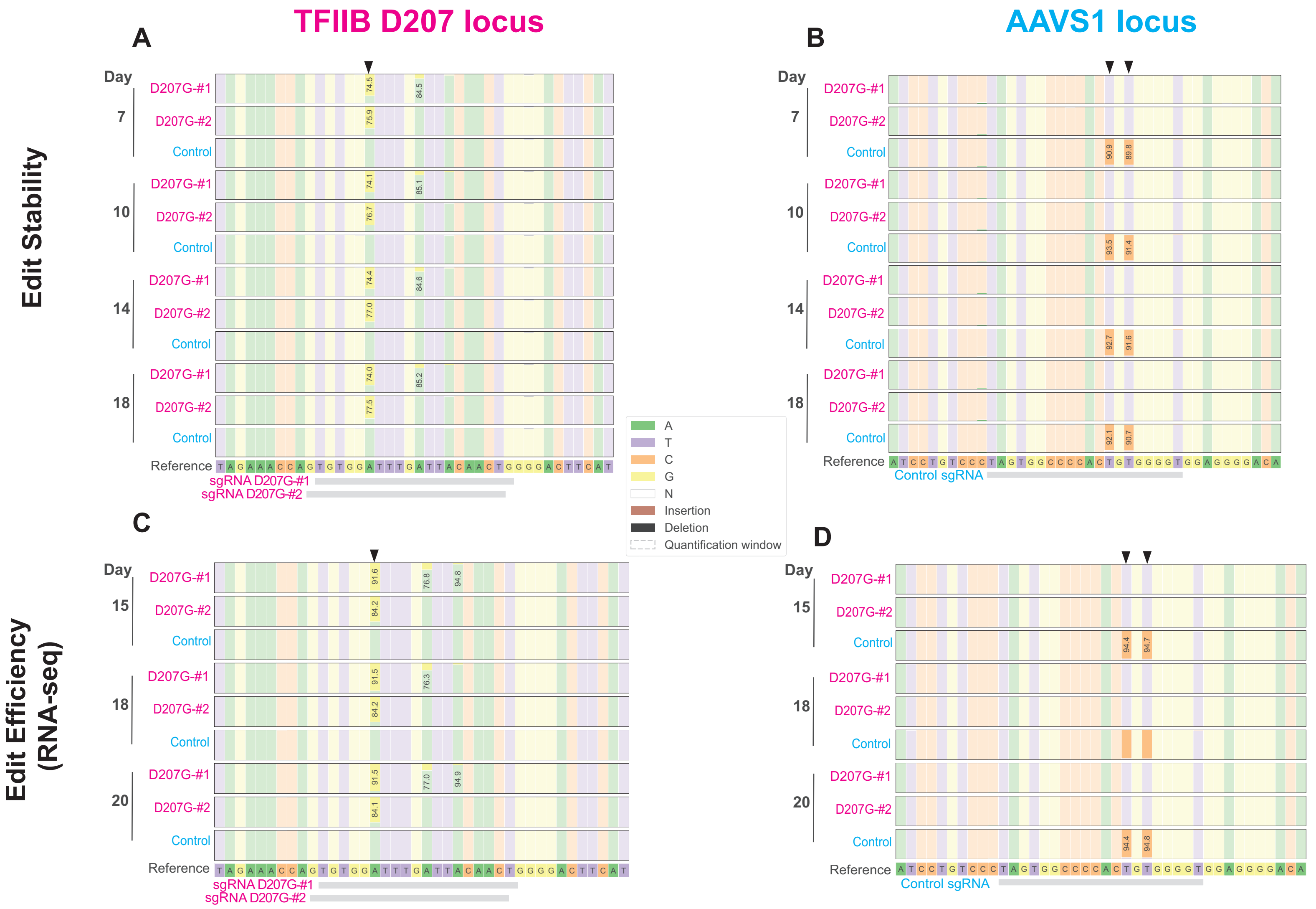


**Supplemental Figure S4. (A-D)** Base editing quantitation at TFIIB and AAVS1 loci from gDNA extracted at the indicated timepoints post-delivery of guide and A•G base editor RNA in Jurkat cells. D207G-#1 and D207G-#2 correspond to sgRNAs targeting the TFIIB D207 locus for editing, and the control population sgRNA targets a dinucleotide motif at the AAVS1 locus. Quantitation was performed using the CRISPResso2 package on PCR amplicon sequencing (Clement et al. 2018; Clement et al. 2019). Black arrows indicate the desired edit loci and sgRNA homology sequences are in gray. Population frequency for bases with 50% < purity < 99% purity are given as percentages for the dominant base. Predictions from the UCSC Genome Browser (Perez et al. 2025) indicated low off-target potentials, primarily in intronic or intergenic regions. **(A-B)** Edit stability was assessed from 7-18 days after editor delivery and did not yield substantial variation at any site. **(C-D)** Editing efficiency in each guide pool was measured over the duration of SLAM-sequencing experiment data collection in main **Fig. 3** to verify sufficient editing. All target edits were present at ~85% or greater in the population. The predominant bystander in D207G-#1 cells had < 15% editing and led only to V209I exchange, a mutation that is unlikely to have consequences for protein function; D207G-#2 cell and control cells had no bystanders.

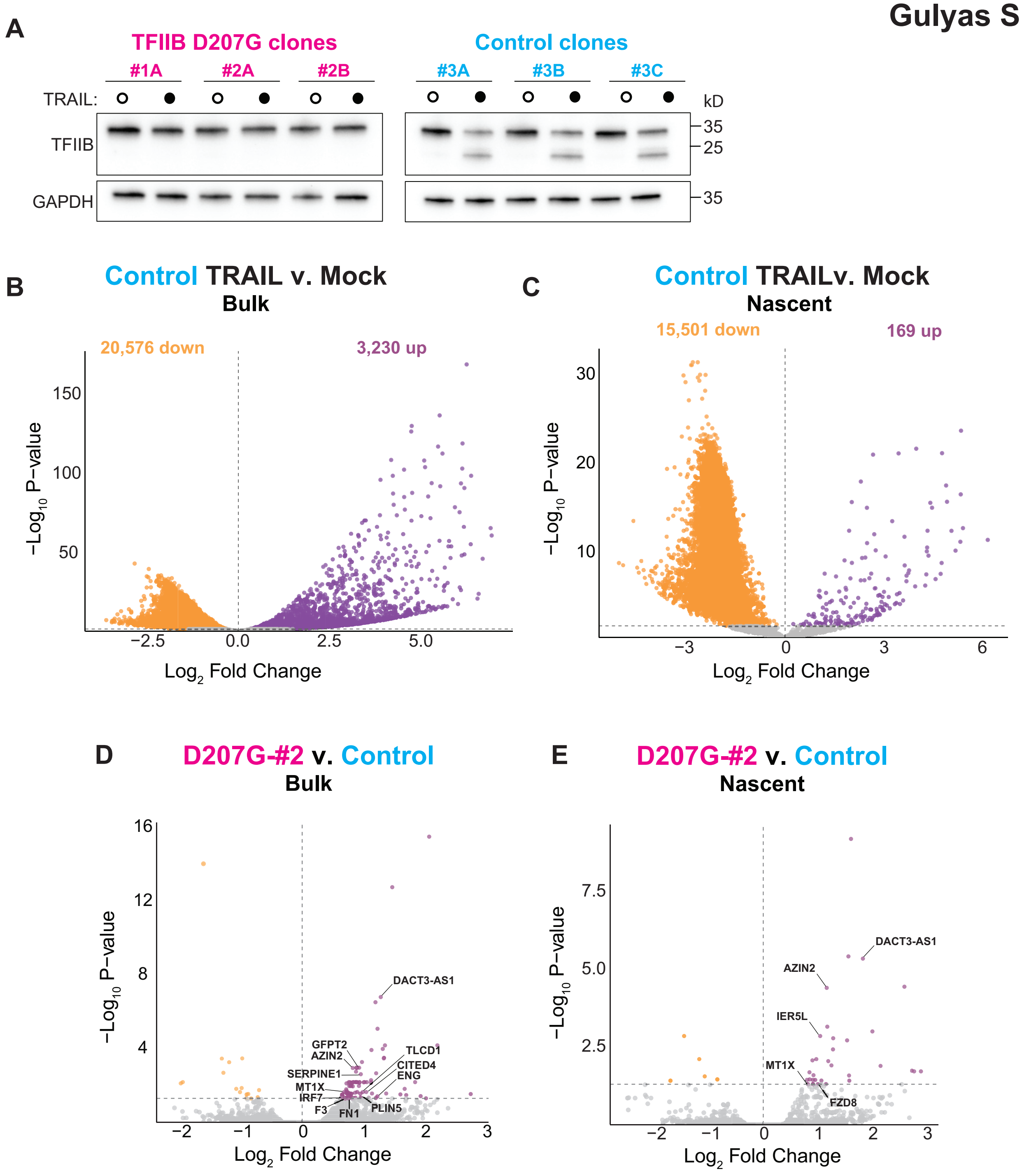


**Supplemental Figure S5. (A)** Western blots of each clonal Jurkat line +/- 250 ng/mL TRAIL ligand for 6 h to confirm total loss of TFIIB cleavage in D207G-edited lines. Lines are labelled using the guide number (#1 and #2 for the corresponding D207G guide, #3 for the control guide) followed by the identity of the clonal line (e.g., A). GAPDH is a loading control. **(B-C)** Bulk *(B)* and nascent *(C)* RNA expression changes from the control edited pooled Jurkat population in main **Fig. 3**. Volcano plots comparing control cells treated with 250 ng/mL TRAIL versus mock diluent for 6 h display significantly upregulated genes under TRAIL conditions in purple and significantly downregulated genes in yellow, confirming bulk and nascent transcriptome repression during apoptosis. **(D-E)** Volcano plots of bulk *(D)* and nascent (*E)* RNA differential gene expression between the TFIIB D207G-#2 guide population and the control population. Significantly downregulated genes are in yellow and significantly upregulated genes are in purple. TRAIL-specific gene upregulations: 56 bulk, 22 nascent.

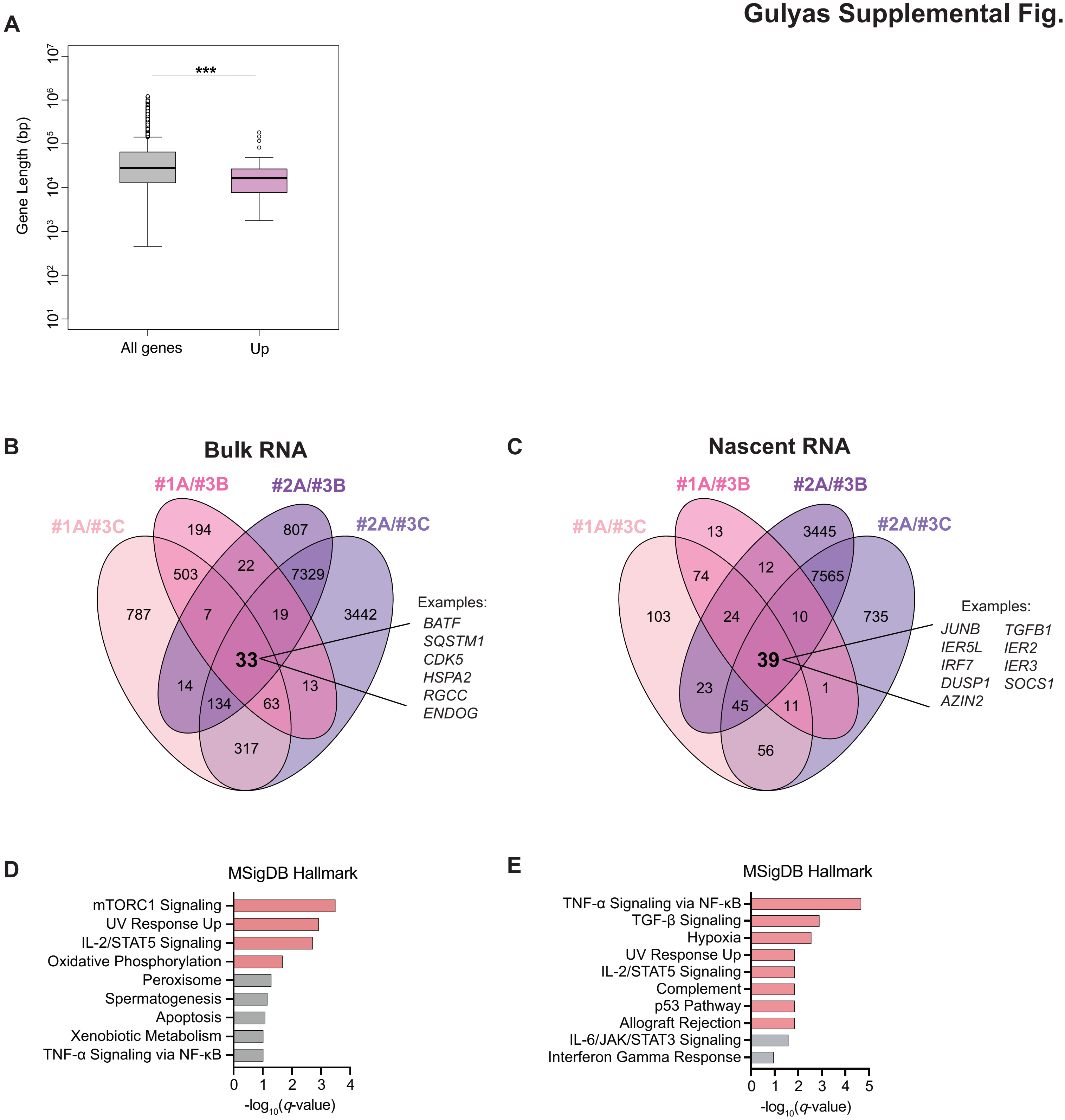


**Supplemental Figure S6. (A)** Boxplot comparing gene lengths for all genes quantifiable in the exon-intron split dataset (n = 10,746) and genes upregulated (purple) in a TRAIL-specific manner in nascent RNA from TFIIB(D207A)-2xStrep expressing BCBL-1 cells relative to wild-type TFIIB-2xStrep control cells. ***$p < 0.001$; Wilcoxon rank sum test with continuity correction. **(B)** Venn diagram of significant TRAIL-upregulated genes from bulk RNA SLAM-seq data in clonal D207G Jurkat cell lines. To stringently compensate for drift, only genes that were consistently upregulated in all pairwise comparisons between each D207G-edited line and control line during TRAIL were considered (33 genes bolded in the center of the diagram); changes also present in mock conditions were discarded. **(C)** As in *B* for nascent RNA. **(D)** MSigDB enrichment for the 33 genes consistently upregulated in bulk RNA (*B*) with terms showing significant *q*-values in pink and non-significant *q*- but significant *p*-values in gray. **(E)** As in *D* for the nascent RNA overlap shown in *C* .

### Gulyas Supplemental Fig. S7

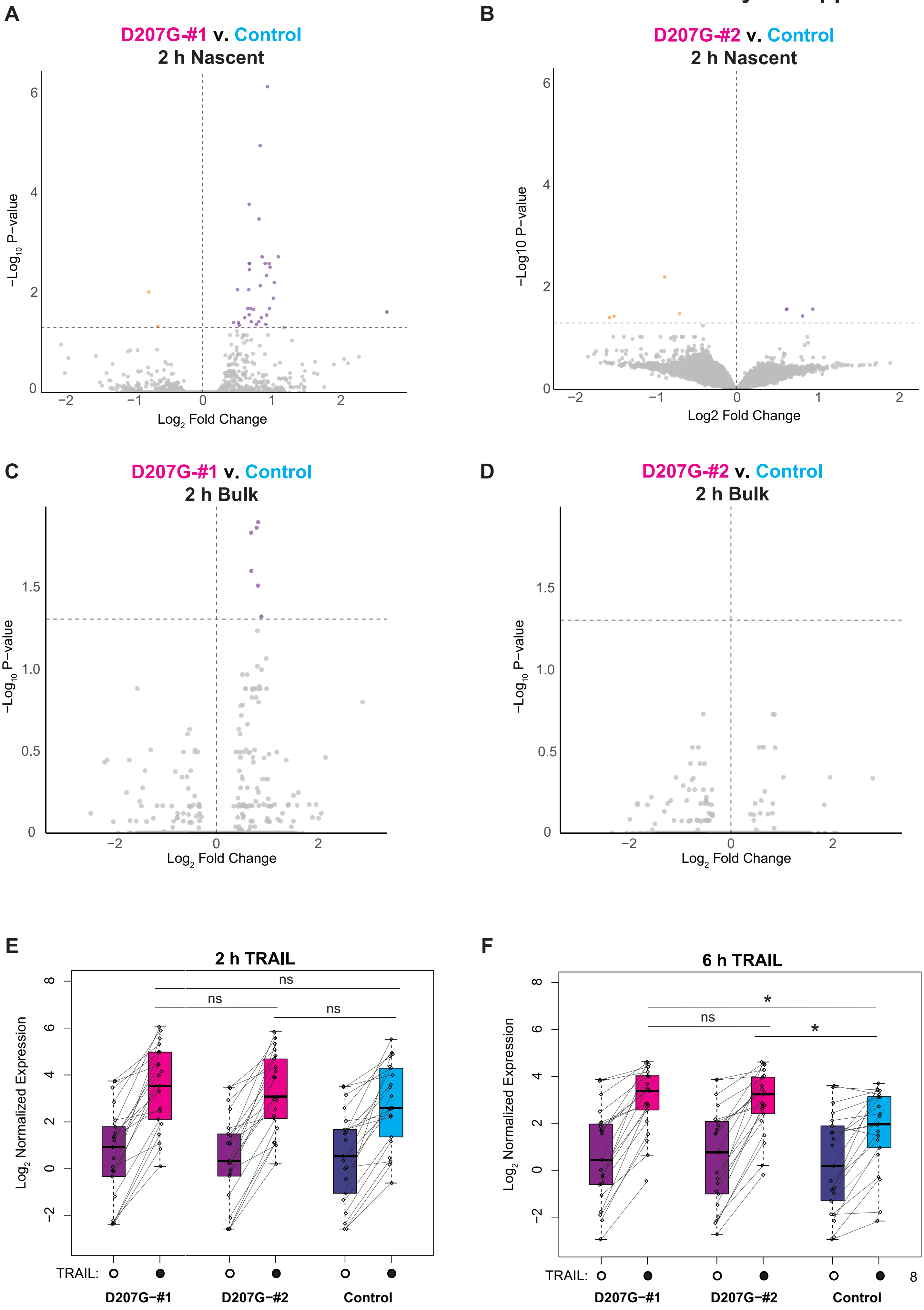

**Supplemental Figure S7. (A-D)** Volcano plots from SLAM-seq comparing D207G-#1 or D207G-#2 Jurkat guide pools to the control pool as indicated after 2 h of 250 ng/mL TRAIL-treatment with 4sU-labelling. *(A-B)* shows changes in nascent RNA expression and (*C-D)* shows bulk. Significantly downregulated genes are in yellow and significantly upregulated genes are in purple. *(A)*: 34 genes up, 2 down; 22/34 genes overlap with loci upregulated in the bulk transcriptome at 6 h post-TRAIL treatment. Minor cleavage begins to occur between 1-2 h post-TRAIL, so significantly upregulated genes in this population are likely highly sensitive to TFIIB loss and beginning to show differential regulation in the D207G-#1 mutant. *(B)* 4 upregulated, 5 downregulated. *(C)* 6 upregulated, no downregulated. *(D)* No significant changes. **(E)** Jittered line and boxplot showing $\log_2$ normalized nascent RNA expression levels after 2 h TRAIL for the 21 loci upregulated in nascent RNA from both D207G-#1 or #2 Jurkat pools (overlapping). **(F)** As in *E*, but for $\log_2$ normalized bulk RNA expression after 6 h TRAIL treatment with 3 h 4sU-labelling. Significance results for E-F are shown for a one-way ANOVA with Tukey HSD: $^*p < 0.05$, ns is nonsignificant.

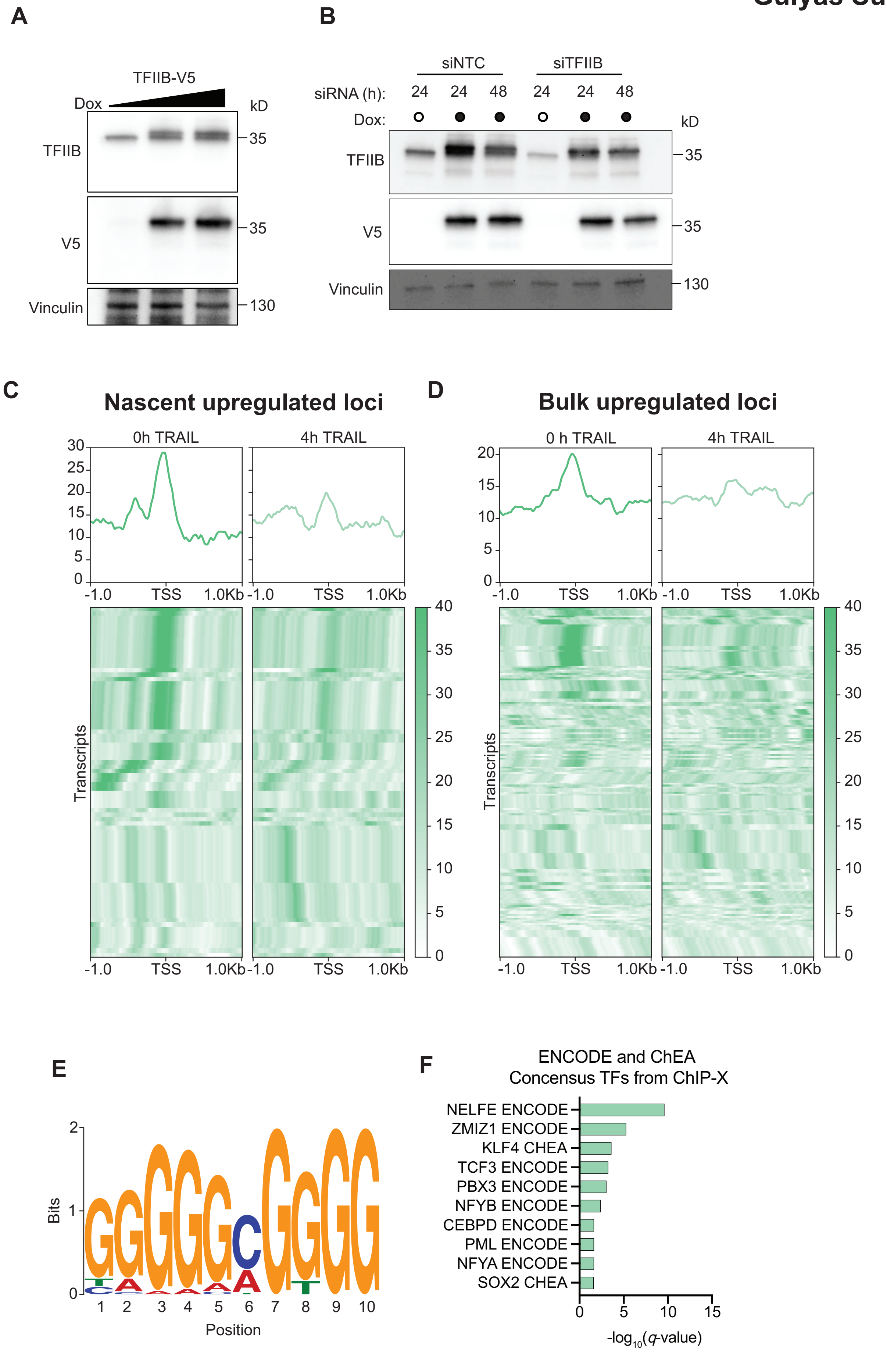
A
TFIIB-V5
Dox
kD
TFIIB
35
V5
35
Vinculin
130
B
siNTC
siTFIIB
siRNA (h):
24
24
48
24
24
48
Dox:
kD
TFIIB
35
V5
35
Vinculin
130
C
Nascent upregulated loci
0h TRAIL
4h TRAIL
30
25
20
15
10
5
0
-1.0
TSS
1.0Kb
Transcripts
40
35
30
25
20
15
10
5
0
D
Bulk upregulated loci
0 h TRAIL
4h TRAIL
20
15
10
5
0
-1.0
TSS
1.0Kb
Transcripts
E
Bits
2
1
0
1
2
3
4
5
6
7
8
9
10
Position
F
ENCODE and ChEA
Concensus TFs from ChIP-X
NELFE ENCODE
ZMIZ1 ENCODE
KLF4 CHEA
TCF3 ENCODE
PBX3 ENCODE
NFYB ENCODE
CEBPD ENCODE
PML ENCODE
NFYA ENCODE
SOX2 CHEA
0
5
10
15
-log10(q-value)

**Supplemental Figure S8. (A-D)** Jurkat lines stably expressing siRNA-resistant, dox-inducible TFIIB-V5 were generated for ChIP-sequencing to monitor TFIIB-V5 promoter occupancy in the absence of endogenous TFIIB. **(A)** Western blot showing the level of TFIIB-V5 relative to endogenous TFIIB following 24h induction with 0, 83, or 166 μg/mL of Dox. Anti-TFIIB antibody recognizes both endogenous and TFIIB-V5 protein; TFIIB-V5 is the upper band appearing in + Dox conditions. Vinculin is a loading control. **(B)** Western blot of Jurkat TFIIB-V5 cells +/- Dox and treated with control (siNTC) or TFIIB-targeting (siTFIIB) siRNA, demonstrating the level of endogenous TFIIB knockdown. **(C)** TFIIB-V5 expression was induced in Jurkats in tandem with endogenous TFIIB siRNA knockdown and TFIIB-V5 occupancy measured by ChIP-sequencing +/- 250 ng/mL TRAIL ligand for 4 h. Metagene heatmaps and profiles for ChIP-seq signal of TFIIB-V5 is shown for the transcript coordinates of genes upregulated in nascent RNA in both D207G-#1 and #2 Jurkat pools when TRAIL-treated relative to the control pool (n = 80). TSS = transcription start site, kilobases plotted up and downstream are indicated. **(D)** As in *C* but for loci upregulated in bulk RNA (n = 195). *C-D* were clustered by detectable RBP1 occupancy to filter out inactive or low signal transcripts and the heatmaps sorted by signal strength. **(E)** MEME analysis of the 500 bp region upstream of the transcription start site for all loci nascently upregulated across TRAIL-treated TFIIB(D207A)-expressing BCBL-1 cells, clonal D207G Jurkat cells, or in the D207G-#1 and D207G-#2 Jurkat pool overlap. A GC-rich motif (E-value = $1.1 \times 10^{-27}$) was present 70/97 sites tested and was discriminative relative to all other detectably expressed genes in the dataset. This corresponds to multiple transcription factors, which included KLF and SP family members. **(F)** Enrichment of transcription factors targeting the genes in *E* was assessed with ENCODE and ChEA Consensus Transcription Factor sites from ChIP-X. All *q*-values are significant.

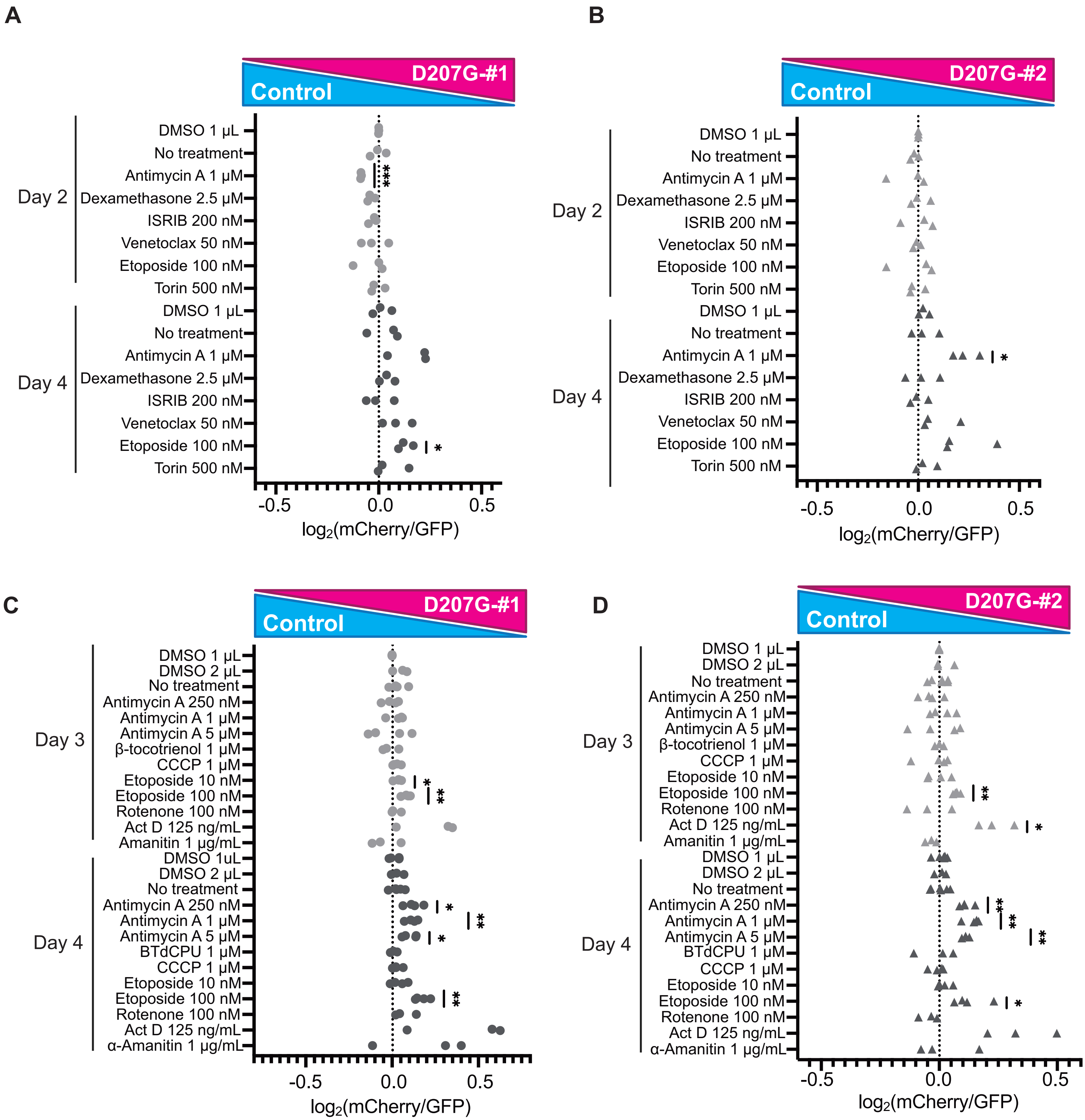


**Supplemental Figure S9.** Initial drug screening for data presented in main **Fig. 5**. Drugs were tested at indicated concentrations and normalized to an appropriate DMSO diluent volume (Abbreviations: Act D- Actinomycin D, CCCP- Carbonyl cyanide m-chlorophenyl hydrazone, ISRIB- integrated stress response inhibitor). **(A-B)** Competition experiments between D207G-#1 (*A*) or D207G-#2 (*B*) guide populations are shown relative to the control; samples were normalized to DMSO timepoints taken at day 2 post-plating to account for variations in initial plating or basal growth. **(C-D)** Full screening results for the data subset presented in main **Fig. 5B**, which includes additional DNA damage agents and mitochondrial stressors. Samples were normalized to DMSO population ratios collected on day 3 post-plating. For *A-D*, $log_2$ of mCherry:GFP ratios for independent replicates are plotted for D207G-#1:control competitions (circles) and D207G-#2:control (triangles) competitions. Light gray indicates an earlier timepoints (day 2-3) and dark gray indicates data collection on day 4. *$p$<0.05, **$p$<0.01, ***$p$<0.001; one sample t-test with theoretical mean of 0.

#### Supplemental Materials and Methods

##### Jurkat pooled line edit validation

After D207G-#1, D207G-#2, or AAVS1 edit pools were recovered and expanded for ≥ 7 days, 1 × $10^6$ cells were harvested for genomic DNA at indicated times using a Monarch Genomic DNA Purification Kit. The TFIIB and AAVS1 target loci were Phusion PCR amplified (NEB) (primers in **Supplemental Table S2**), and product was purified and concentrated with a Monarch PCR & DNA Cleanup Kit. Amplicons were prepared and sequenced on an Illumina MiSeq instrument (2 × 300 bp -v3 reagents) at the Innovative Genomics Institute NGS facility (UC Berkeley). Amplicon sequencing was visualized and editing efficiency assessed using the CRISPResso2 package (Clement et al. 2019).

##### Jurkat clonal line generation and edit validation

Clonal lines were derived by single-cell dilution cloning 48 h post-delivery of the base editor and guide nucleofection into Jurkats. Briefly, cells were diluted to ~ 1 cell/mL and 100 μL per well was plated in 4 × 96 well plates per guide with an additional 100 μL of fresh media. Cells were then recovered and clonal populations expanded and frozen along with gDNA extraction as described above. PCR amplicons at the edit loci were submitted for long-read sequencing through Plasmidsaurus. Pristine editing was manually confirmed by analyzing the sequence traces in Snapgene.

##### ChIP-sequencing system validation and experimentation

*Cell line generation*

6xHis-TFIIB-V5 expressing Jurkat cells were generated with the Neon Transfection System (Invitrogen) according to the manufacturer's instructions with the following parameters: 2.5 μg vector DNA and 0.125 μg transposase plasmid was nucleofected (1700 V, 1 × 20 ms pulse-length) into 5 × $10^6$ low-passage cells in 100 μL tip with resuspension buffer T; 10 μg/mL blasticidin HCl (Gibco) selection commenced 24 h later.

*siRNA knockdown for validation and ChIP-sequencing samples*

After 24 h of doxycycline-induction (2.5 μg/mL) of siRNA-resistant 6xHis-TFIIB-V5 in Jurkat cells, endogenous protein was knocked down by siRNA. 3 × $10^6$ Jurkat cells were prepared in 100 μL of resuspension buffer T (Invitrogen) and 3 μM of the following Dharmacon siRNAs: On-Target Plus Non-targeting control pool and Accell Non-targeting control pool for non-targeting control cells, or On-Target Plus human *GTF2B* siRNA SmartPool and Accell human *GTF2B* siRNA SmartPool for TFIIB knockdown cells. Cells were nucleofected (1700 V, 1 × 20 ms pulse-length) with the Neon Transfection System (Invitrogen) according to the manufacturer's instructions. siRNA treatments for ChIP-sequencing were performed identically on 10 × $10^6$ Jurkat cells with 10 μM each of On-Target Plus human *GTF2B* siRNA SmartPool and Accell human *GTF2B* siRNA SmartPool.

*Library preparation and sequencing*
Jurkat cells were prepared and siRNA-treated as described. After 24 h, 40 × $10^6$ cells were dosed with 250 ng/mL SUPERKILLER TRAIL (Enzo) or mock diluent for 4 h. Samples were prepared and DNA sequenced as described previously (Lari et al. 2025) with the following modifications: during immunoprecipitation, 10 µg of purified chromatin was incubated with 5 µg rabbit monoclonal anti-RPB1 NTD (clone D8L4Y; Cell Signaling 14958), rabbit polyclonal anti-V5 (Abcam ab15828), or rabbit IgG (Abcam ab37415) overnight. During library preparation, yeast spike-in DNA (Cell Signaling) was added for downstream normalization.

*Data analysis and visualization*
Sequencing quality was assessed with FastQC, and files were trimmed with Trimmomatic v. 0.39 (Bolger et al. 2014). The human (hg38) and yeast (SacCer3) genomes were manually indexed using the bowtie2-build command. Processed sequencing files were aligned to the genomes with bowtie2 v. 2.5.2. Calculated spike-in normalization factors were proportional to the number of spike-in reads per sample and applied to generate BigWig files with bamCoverage (--binSize 10, --smoothLength 30, --extendReads, --centerReads, --normalizeUsing None, and --scaleFactor with the respective spike-in normalized scale factor). Biological duplicate files were averaged with the deepTools bigwigCompare function.

Matrices for V5 and RPB1 (available at GSE302742) coverage profiles were made with deepTools computeMatrix reference-point command on the averaged BigWig files (-S), with --referencePoint TSS, --skipZeros, --missingDataAsZero. Bedfiles were obtained from Ensembl BioMart transcript coordinates. V5 ChIP-seq signal for only transcripts with detectable RPB1 occupancy in either condition was plotted with plotHeatmap as a histogram with 10bp bins, 100 bins/gene and sorted by signal strength. ChIP-seq data are deposited on the GEO database (GSE302742).

#### MEME analysis

MEME (Bailey et al. 2015) was used to identify discriminatory motifs in the upstream regions of nascently upregulated genes in TFIIB(D207A)-expressing BCBL-1 cells, clonal D207G Jurkat cells, or those overlapping in D207G-#1 and D207G-#2 Jurkat pools relative to their respective controls. All other detectably expressed genes in the dataset were used as a negative control. Input data was generated with the hg38 genome FASTA file by obtaining the DNA sequence 500 bp upstream of the TSSs for all genes analyzed (n = 97). All settings used for MEME were set to defaults with a motif width of 10. The Tomtom motif comparison tool (Gupta et al. 2007; Bailey et al. 2015) identified corresponding transcription factors.
